# The maize Aliphatic Suberin Feruloyl Transferase genes affect leaf water movement but are dispensable for bundle sheath CO_2_ concentration

**DOI:** 10.1101/2020.09.30.321265

**Authors:** Rachel A. Mertz, Patrick Ellsworth, Patricia Ellsworth, S. Lori Tausta, Susanne von Caemmerer, R. Howard Berg, Timothy Nelson, Nicholas C. Carpita, Thomas P. Brutnell, Asaph B. Cousins

## Abstract

C_4_ grasses often outperform C_3_ species under hot, arid conditions due to superior water and nitrogen use efficiencies and lower rates of photorespiration. A method of concentrating CO_2_ around the site of carbon fixation in the bundle sheath (BS) is required to realize these gains. In NADP-malic enzyme (NADP-ME)-type C_4_ grasses such as maize, suberin deposition in the BS cell wall is hypothesized to act as a diffusion barrier to CO_2_ escape and O_2_ entry from surrounding mesophyll cells. Suberin is a heteropolyester comprised of acyl-lipid-derived aliphatic and phenylpropanoid-derived aromatic components. To disrupt BS suberization, we mutated two paralogously duplicated, unlinked maize orthologues of Arabidopsis thaliana *ALIPHATIC SUBERIN FERULOYL TRANSFERASE, ZmAsft1* and *ZmAsft2*, using closely linked *Dissociation* transposons. Loss-of-function double mutants revealed a 97% reduction in suberin-specific omega-hydroxy fatty acids without a stoichiometric decrease in ferulic acid. However, BS suberin lamellae were deficient in electron opaque material, and cohesion between the suberin lamellae and polysaccharide cell walls was attenuated in double mutants. There were no other morphological phenotypes under ambient conditions. Furthermore, there was no significant effect on net CO_2_ assimilation at any intercellular CO_2_ concentration, and no effect on ^13^C isotope discrimination relative to wild type. Thus, *ZmAsft* expression is not required to establish a functional CO_2_ concentrating mechanism in in maize. Double mutant leaves exhibit elevated cell wall elasticity, transpirational, and stomatal conductance relative to WT. Thus, the *ZmAsft* genes are dispensable for gas exchange barrier function but may be involved in regulation of leaf water movement.

**One-sentence Summary:** Double mutants of two paralogously duplicated maize Aliphatic Suberin Feruloyl Transferase (ZmAsft) genes exhibit reduced aliphatic suberin content, cell wall cohesion defects, and elevated leaf transpiration, but no changes in CO2 assimilation relative to wild type.

## INTRODUCTION

Cereal crops are among our most productive sources of food, feed, and fuel. Just three staple cereals, rice (*Oryza sativa*), wheat (*Triticum aesetivum*), and maize (*Zea mays*) are estimated to provide 42.5% of dietary calories consumed directly by humans and 30% of dietary protein (FAO, 2015). Global food demand is forecasted to increase significantly over the coming decades and will be double the current level by 2050 (Tillman *et al.*, 2011). However, at their current rate of growth, maize, rice, and wheat yields are forecasted to increase by only 67%, 42%, and 38%, respectively, far short of the level required to keep pace with population growth (Ray *et al.*, 2013). It was estimated that a 50% yield increase on existing acreage without additional fertilizer inputs could be generated in the C_3_ cereal crop rice by engineering novel cultivars utilizing the NADP-ME-type C_4_ photosynthetic pathway (Mitchell and Sheehy, 2006; Langdale, 2011; von Caemmerer *et al.*, 2012).

In grasses utilizing C_4_ photosynthesis, CO_2_ is incorporated into a 4-carbon organic acid in the leaf mesophyll (M) cells by phosphoenol pyruvate carboxylase (PEPC) and released proximal to the active site of ribulose-1,5-*bis*-phosphate carboxylase/oxygenase (Rubisco) in an internal cell layer, the bundle sheath (BS), located adjacent to the leaf vasculature (termed Kranz Anatomy; reviewed in Sage, 2004). C_4_ species are broadly grouped into three classes, termed NAD-ME, PEPCK, and NADP-ME-type, based on the primary decarboxylase used to release CO_2_ within the BS (reviewed in Furbank, 2011). This CO_2_ concentrating mechanism (CCM) protects the plant from photorespiration under hot, arid conditions and significantly increases the water and nitrogen use efficiencies relative to C_3_ species (Sage, 2004; Taylor *et al.*, 2010). Establishment of an efficient CCM requires a low ratio of the CO_2_ leak rate out of the BS relative to the rate of PEP carboxylation (leakiness; *ϕ*). High *ϕ* is typically associated with low efficiency of the CCM because concentrating CO_2_ in the BS requires energy through the consumption of two ATP in PEP regeneration. However, diffusivity across the bundle sheath is a balance between retaining CO_2_ in the bundle sheath (low *ϕ*), permitting O_2_ diffusion out, and allowing ample metabolite diffusion out to support maximum rates of CO_2_ fixation. Thus, in addition to cell-type specific expression of key carbon concentrating genes, multiple anatomical modifications may be required to engineer an efficient C_4_ CCM into C_3_ species, including greater vein density, increased plasmodesmatal connections between BS and M, and the modification of the cell walls at the BS/M interface (reviewed in Nelson, 2011).

Cell wall attributes have long been suspected to influence CO_2_ diffusion across the BS/M interface because maintaining high resistance to CO_2_ diffusion is essential for concentrating CO_2_ within the BS. Over forty years ago, suberin lamellae (SL) were proposed to act as gas diffusion barriers, facilitating CO_2_ concentration within the BS while blocking penetration of atmospheric O_2_, thereby reducing photorespiration (Laetsch, 1971). Suberized BS cells are not uniformly distributed across the grasses. Although they are generally present in NADP-ME and PEPCK-type C_4_ species, both NAD-ME-type C_4_ and C_3_ grasses have an unsuberized parenchymatous BS (Hattersley & Browning, 1981; Eastman *et al.*, 1988a). Carbon isotope discrimination ratios and estimates of *ϕ* are comparable between suberized and unsuberized C_4_ grasses (Henderson *et al.*, 1992; von Caemmerer *et al.*, 2014; Kromdijk *et al.*, 2014). This implies either that the lignocellulosic walls and elongated symplasmic diffusion path of NAD-ME species constitutes an equally effective gas exchange barrier (Hattersley and Browning, 1981), or that SL are not essential for the CCM. However, cell wall thickness in itself does not appear to be important in increasing resistance to CO_2_ diffusion (Evans and Von Caemmerer, 1996, Leegood, 2002, von Caemmerer and Furbank, 2003). Thus, the function of the SL must be tested before engineering these structures into C_4_ rice.

In addition to its proposed role in gas exchange, BS suberization also creates barriers to transpiration by restricting the apoplastic diffusion path of water and solutes across the BS/M interface (O’Brien and Carr, 1970). For example, solutes are blocked by the SL but diffuse through the unsuberized radial walls between adjoining BS cells, which represent a contiguous apoplastic path out of the vasculature (Evert *et al.*, 1985; Botha and Evert, 1986). Thus, suberization may increase resistance and reduce hyrdraulic conductivity outside the xylem (K_ox_). The quantity, composition, and ultrastructure of the SL all contribute to permeability, though the relative contributions of each remain to be elucidated, particularly for the BS (Schreiber, 2010; Mertz and Brutnell, 2014).

Suberin is a heteropolyester comprised of both aliphatic and aromatic components. In maize, the principal aliphatic monomers are C_16_-C_28_ fatty acids and several classes of oxidized and reduced derivatives, including primary alcohols, 2-hydroxy fatty acids, omega-hydroxy fatty acids, and alpha, omega-dicarboxylic acids (Supplemental Figure S1A; Espelie and Kolattukudy, 1979; Zeier *et al.*, 1999). The same aliphatic monomer classes are present in the epidermal cuticle (Espelie and Kolattukudy, 1979); however, suberin is consistently enriched in very long chain (≥ C_20_) aliphatic species and aromatic monomers relative to cutin across multiple taxa (Pollard *et al.*, 2008). The predominant aromatic monomers are the hydroxycinnamic acids *p*-coumaric and ferulic acid (Zeier *et al.*, 1999). Hydroxycinnamic acids are not specifically associated with suberin in maize leaves. Maize has a Type II primary cell wall with extensive esterification of ferulic acid to the arabinosyl side chains of the hemicellulose glucuronoarabinoxylan (GAX; reviewed in Carpita and Gibeaut, 1993). Likewise, substantial quantities of *p*-coumaric acid are present in maize lignin as esterified side groups of sinapyl alcohol residues (Hatfield *et al.*, 2008). Although the suberin, GAX, and lignin-related hydroxycinnamoyl transferases are thought to be phylogenetically distinct enzymes belonging to the BAHD acyltransferase super-family (Mitchell *et al.*, 2007; Bartley *et al.*, 2013), no aliphatic suberin O-hydroxycinnamoyl transferases have been characterized to date in any monocot.

Recently, multiple hydroxycinnamoyl transferases involved in cutin, suberin, and suberin-associated wax biosynthesis have been characterized in the model dicots Arabidopsis (*Arabidopsis thaliana*), potato (*Solanum tuberosum*) and poplar (*Populus trichocarpa*; reviewed in Molina and Kosma, 2015). Arabidopsis *ALIPHATIC SUBERIN FEULOYL TRANSFERASE* (*AtASFT*; Gou *et al.*, 2009; Molina *et al.* 2009) and potato *ω-Hydroxy fatty acid/fatty alcohol hydroxycinnamoyl transferase* (*StFHT*; Serra *et al.*, 2010) are required for normal incorporation of ferulic acid and hydroxylated acyl lipids into the suberin polyester in an apparent 1:1 stoichiometry (Supplemental Figure S1B; Molina et al., 2009). Expression of *AtASFT* and *StFHT* is dispensable for normal suberin ultrastructure but required to form functional barriers to the unrestricted diffusion of salts and water across the seed coat and tuber periderm, respectively (Gou et al., 2009; Molina et al., 2009; Serra et al., 2010). Conversely, mutants of the cutin feruloyl transferase *DEFICIENT IN CUTIN FERULATE (AtDCF)* and the wax-related *FATTY ALCOHOL:CAFFEOYL-CoA CAFFEOYL TRANSFERASE (AtFACT)* have negligible effects on barrier function (Rautengarten et al., 2012; Kosma et al., 2012). Thus, the maize *ASFT* homologues are promising targets with which to disrupt BS suberization without significantly compromising cutin barrier function.

Previously, we delineated a putative suberin biosynthetic pathway and identified a subset of genes expressed concurrently with vascular sheath suberization in both maize and rice (Li *et al.*, 2010; Wang *et al.*, 2014). Of the candidate genes in this subset, only the *ZmAsft* genes had one or more *Dissociation* transposons from the maize *Activator/Dissociation* collection in sufficiently tight genetic linkage to generate targeted insertion mutants (Supplemental Table S1; Ahern *et al.*, 2009; Vollbrecht *et al.*, 2010). To investigate the role that the ultrastructure of suberin plays in the bundle sheath cells, we generated an allelic series of mutations in *ZmAsft1* and *ZmAsft2* by targeted insertional mutagenesis using *Dissociation* elements. *ZmAsft1* and *ZmAsft2* were found to be redundantly essential for normal accumulation of very long chain omega-hydroxy fatty acids in leaf polyesters. Furthermore, the ultrastructure of the BS cell wall, but not the epidermal cuticle, was compromised in double mutants. We tested the effect of the double mutation on gas diffusion, including O_2_ diffusion under waterlogging and CO_2_ diffusion through the bundle sheath. We also evaluated the effect of the double mutation on water transport through the transpiration stream.

## RESULTS

### *ZmAsft1* and *ZmAsft2* are expressed concurrently with suberin synthesis in maize leaves

*ZmAsft1* (*ZmHydroxycinnamoyltransferase10*; Zm00001d012274) and *ZmAsft2* (Zm00001d042723) are syntenic paralogues resulting from an ancestral polyploidy event in the maize lineage after it diverged from *Sorghum* (Supplemental Figure S2; Schnable *et al.*, 2009). *ZmAsft3* (Zm00001d024864) is an unlinked paralogue of *ZmAsft2* (Supplemental Figure S2). Although the protein encoded by the predicted *ZmAsft3* gene model contains the HXXXD and DFGWG motifs of a canonical BAHD acyltransferase (D’Auria, 2006), it lacks the N-terminal amino acids encoded by Exons 1 and 2 of *ZmAsft1* and *ZmAsft2*, and is not transcribed in either B73 or W22 leaves (Supplemental Figure S3, A and B). Thus, we concluded that *Asft3* is likely a pseudogene and *ZmZmAsft1* and *ZmZmAsft2* were selected for further characterization.

*ZmAsft1* and *ZmAsft2* are expressed in multiple vegetative tissues that undergo suberization, including primary roots and developing leaves (Supplemental Figures S4A and S4C; Stelpflug *et al.*, 2015). In developing juvenile leaves of the inbred B73, expression is restricted to a narrow spatiotemporal region concurrent with maturation of metaxylem vessels (Li *et al.*, 2009; Wang *et al.*, 2014). To determine whether *ZmAsft1* and *ZmAsft2* are expressed similarly in the W22 inbred background, we performed qRT-PCR. As in B73, both paralogues were strongly expressed in the transition zone proximal to the point of leaf emergence, with negligible expression at the leaf base or in emerging source tissue (Fig. 1A). Thus *ZmAsft1* and *ZmAsft2* are likely to play similar roles during leaf maturation in both genetic backgrounds.

**Figure 1.**
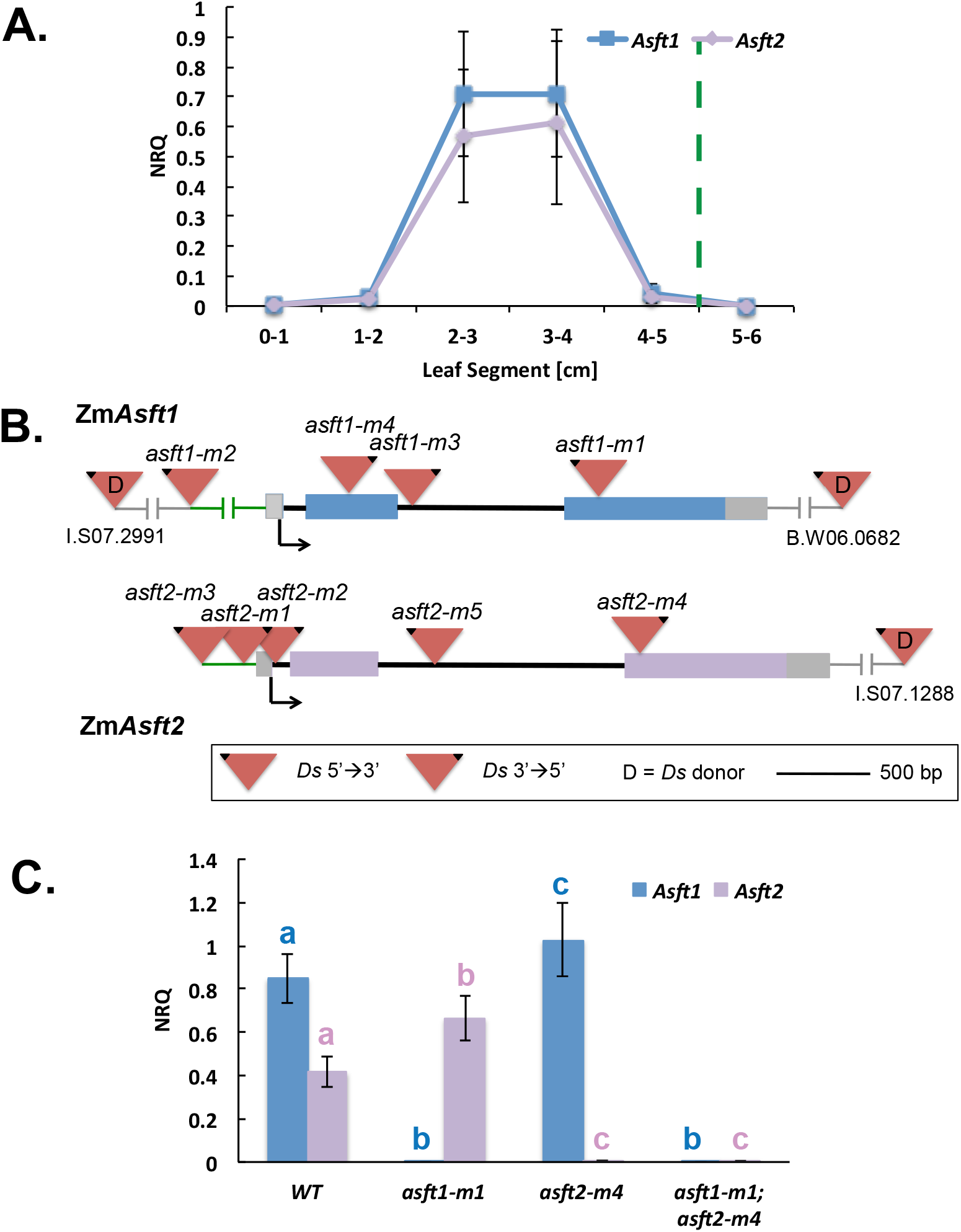
Identification and mutagenesis of two putative maize *Aliphatic Suberin Feruloyl Transferase (ZmAsft)* genes. **A.** Developing third leaf expression profile of *ZmAsft1* and *ZmAsft2* divided into 1 cm increments relative to the leaf base. The point of emergence is denoted by a dashed green line. Data are presented as means plus standard deviations of normalized relative quantities (NRQ) of 3 biological replicates. **B.** Mutagenesis of the *ZmAsft* genes using *Dissociation (Ds)* transposons. Triangles denote *Ds* transposons; the black corner indicates the 5’ end. Blue, purple and gray boxes denote exons and UTRs of *ZmAsft1* and *ZmAsft2*, respectively. **C.** qPCR analysis of *Zmasft* mutants in the region of maximal expression denoted in Figure 1A. Letters indicate significant differences between genotypes by a Tukey-Kramer post-hoc test (*p* < 0.05) following a significant one-way ANOVA (*p* < 0.001).

Although feruloylation of Arabidopsis suberin and cutin are performed by two separate but closely related acyltransferases, phylogenetic analysis of the ZmASFT sub-clade did not reveal a clear maize orthologue of AtDCF (Fig. 1A). To investigate whether the *ZmAsft* genes are expressed in both suberized and cutinized leaf tissues, we performed laser capture microdissection and RT-PCR of bundle sheath and epidermal cells from the region of maximal expression defined in Figure 1A. *ZmAsft1* and *ZmAsft2* are expressed in both bundle sheath and epidermal cells (Supplemental Figure S4B). Therefore, *ZmAsft1* and *ZmAsft2* are each likely to contribute to both cutin and suberin biosynthesis. Thus, both paralogues were mutagenized by targeted reverse genetics using *Dissociation* transposons.

### The *ZmAsft* genes are amenable to mutagenesis with *Dissociation* transposons

The high degree of sequence similarity (94% identity of coding sequence; Supplemental Figure S2A) and similar expression patterns indicated that *ZmAsft1* and *ZmAsft2* were likely functionally redundant. Thus, *Dissociation* (*Ds*) transposon donors in tight linkage to *ZmAsft1* (I.S07.2991; 71.9 kb upstream) and *ZmAsft2* (I.S07.1288; 29.3 kb downstream) were selected from the *Activator/Dissociation* (*Ac/Ds*) collection and targeted reverse genetic screens were carried out for both paralogues.

A single insertion allele was recovered within Exon 3 of *ZmAsft1* (*asft1-m1::Ds*; Fig. 1B; Supplemental Table S1). The low insertion frequency (0.019%) prompted the selection of a more tightly linked *Ds* donor, B.W06.0682 (68.8 kb downstream of *ZmAsft1*). Three additional insertion alleles, *asft1-m2::Ds, asft1-m3::Ds*, and *asft1-m4::Ds* were identified (0.17%; Fig. 1B). We confirmed that *asft1-m1::Ds* was a null allele by qRT-PCR (one-way ANOVA *p* < 0.001; Fig. 1C). The Exon 2 insertion event *asft1-m4::Ds* also significantly attenuated transcript accumulation (one-way ANOVA *p* < 0.001; Supplemental Figure S5A) and resulted in a premature stop codon within the *Ds* transposon (data not shown). As the insertion occurred upstream of the HXXXD and DFGWG motifs, we concluded that any residual transcripts were likely to encode a non-functional protein.

Despite tight linkage between *ZmAsft2* and donor I.S08.1288, only three insertion alleles (*asft2-m1::Ds, asft2-m2::Ds*, and *asft2-m3::Ds*) were recovered (0.082 %; Fig. 1B and Supplemental Table S1). Furthermore, neither *asft2-m2::Ds* nor *asft2-m3::Ds* were null alleles (Supplemental Figure S5B). An intragenic remobilization of *asft2-m2::Ds* was performed to generate two stronger alleles, *asft2-m4::Ds* in Exon 3 and *asft2-m5::Ds* in Intron 2, each of which retained a duplicate copy of *asft2-m2::Ds* in Intron 1 (0.22%; Fig. 1B and C; Supplemental Figure S5A).

The single mutants *asft1-m1::Ds* and *asft1-m4::Ds* were crossed to *asft2-m4::Ds* and *asft2-m5::Ds* to generate double mutants. There were no obvious morphological differences in any allelic combination of double mutants compared to their isogenic WT siblings under ambient conditions, as was previously reported for *AtASFT* (Molina *et al.*, 2009). Thus, chemical profiling of cell wall polyesters was necessary to differentiate suberin mutants from wild type siblings.

### The *ZmAsft* genes are redundantly essential for aliphatic suberin deposition in leaf and root polyesters

Suberized regions of ten-day-old third leaf laminae were harvested from *asft1-m1*; *asft2-m4* single and double mutants and isogenic WT siblings, and the tissue was mechanically fractionated to isolate BS strands. Although this preparation enriched for suberized BS strands relative to total leaf tissue, some cutinized pavement and guard cells of the epidermis were retained in the samples. The majority of the monomers identified following base-catalyzed transmethylation and silylation of extractive-free cell walls were hydroxycinnamic acids (HCA) in both mutant and WT samples (Fig. 2A). Among aliphatic monomers, the distribution of primary alcohols, fatty acids, omega-hydroxy fatty acids (ω-OH FA), α,ω-dicarboxylic acids, and poly-hydroxy fatty acids were broadly similar between all four genotypes (Fig. 2A). However, when the individual ω-OH FA species were partitioned by chain length, C_22:0_-C_30:0_ very long chain omega-hydroxy fatty acids (ω-OH VLCFA) were strongly and specifically attenuated in double mutants, whereas C_16:0_-C_18:0_ and C_18:1_ long chain species (ω-OH LCFA) were unaffected (one-way ANOVA, *p* < 0.001 for all ω-OH VLCFA species; Fig. 2B). Similar reductions in ω-OH VLCFA were observed in fully expanded third leaves of *asft1-m4; asft2-m5* double mutants (Supplemental Figure S6, A and B) and in *asft1-m1; asft2-m4* double mutant primary roots (Supplemental Figure S7). Similar results were also observed for samples containing purified bundle sheath strands devoid of epidermal cutin (Supplemental Figure S8). Thus, *ZmAsft1* and *ZmAsft2* are redundantly essential for normal accumulation of C_22:0_-C_30:0_ ω-OH VLCFA in both leaf and root suberin polyesters.

**Figure 2.**
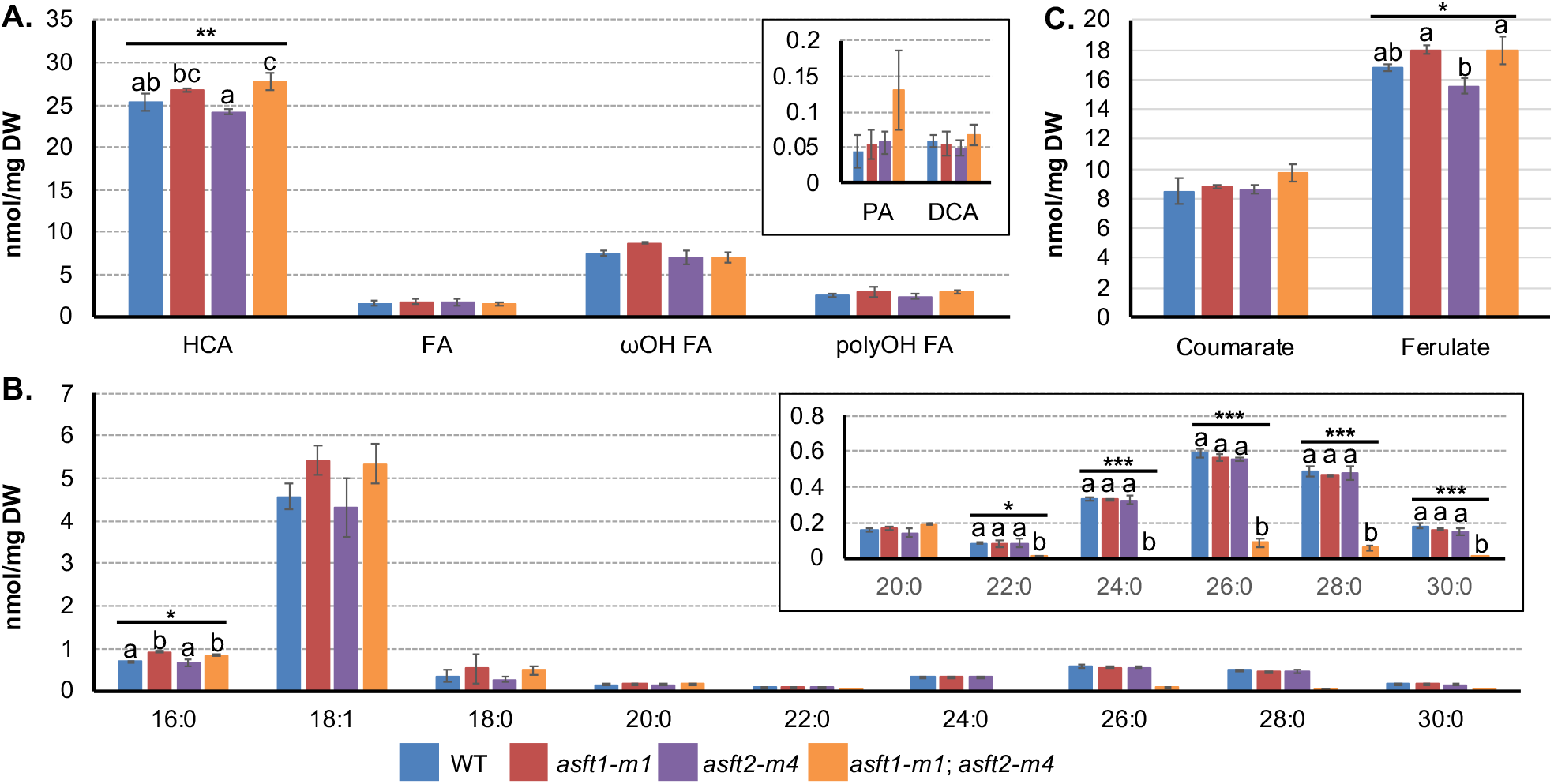
Polyester monomers from isolated bundle sheath strands. **A.** Overview of the major monomer classes. HCA, hydroxycinnamic acids; FA, fatty acids; ωOH FA, omega-hydroxy fatty acids, polyOH FA, poly-hydroxy fatty acids; PA, primary alcohols; DCA, α,ω-dicarboxylic acids. Letters denote significant differences between genotypes determined by a Tukey-Kramer post-hoc test (*p* < 0.05) following a significant one-way ANOVA (*p* < 0.05). Values are averages with standard deviations of three biological replicates. **B.** Chain length distributions of ωOH FA. Inset: Higher resolution image of C_20:0_-C_30:0_ very long chain monomers. **C.** HCA monomers. Letters as defined in Figure 2A.

In dicot *asft* and *fht* mutants, reduced aliphatic monomer levels coincided with a 50-90% decrease in total ferulic acid content (Gou *et al.*, 2009; Molina *et al.*, 2009; Serra *et al.*, 2010). Because maize has a type II primary cell wall with substantial ferulic acid esterification to the arabinose side chains of glucuronoarabinoxylan (GAX; Kato and Nevins, 1985), we expected to observe a smaller total ferulate reduction in maize double mutants relative to the model dicots. However, we observed no stoichiometric decrease in the HCAs ferulic, *p*-coumaric, or caffeic acid concurrent with the reduction in ω-OH VLCFA in either leaves or roots of the double mutant relative to WT (Fig. 2C; Supplemental Figures S6C, S7C, S8C). We investigated whether increased GAX feruloyation masked a ferulate deficiency in the double mutant polyesters. Dilute acid was utilized to selectively hydrolyze 5-O-hydroxycinnamoyl-L-arabinose side chains from the xylan backbone of GAX, which were quantified by UV-HPLC (Supplemental Figure S9A & S9B; Saulnier *et al.*, 1995; Bartley *et al.*, 2013). Both feruloyl and coumaroyl-arabinose dimers were identified, but neither differed significantly between double mutant and WT (Student’s t-test *p* ≥ 0.05; Supplemental Figure S9C and S9D). Likewise, there was no effect on arabinose side chain abundance or xylan branching as determined by a linkage-methylation analysis (Supplemental Figure S9E). Taken together, these data provide no evidence for significant re-allocation of HCAs from polyesters onto GAX in double mutants.

### Bundle sheath suberin ultrastructure is compromised in *Zmasft* double mutants

During the isolation of bundle sheath strands for chemical analyses, *asft1-m1; asft2-m4* double mutant samples appeared lighter in color than single mutant or WT siblings following homogenization (Supplemental Figure S10A). When the preparations were compared under a light microscope, WT and single mutant samples were comprised of isolated BS strands with predominately intact cells containing abundant chloroplasts. Conversely, nearly all of the cells of double mutant BS strands were broken along the outer tangential walls and devoid of plastids (Supplemental Figure S10B). Thus, we observed the ultrastructure of the suberin lamellae (SL) by transmission electron microscopy. In WT and both single mutants, the SL had a classical “tramline” architecture comprised of a central electron lucent region bounded by two electron opaque bands at the interface of the SL with the primary and tertiary cell walls (Fig. 3A). Conversely, the double mutant SL appeared entirely electron lucent at all cell wall positions, and the osmiophilic tramlines delimiting WT lamellae were abolished (Fig. 3A). This phenotype was completely penetrant and affected all BS cells in the minor veins that we observed. The SL was thickened and polylamellate in the vicinity of plasmodesmata in all genotypes (Fig. 3B). However, although the double mutant SL thickened in the presence of plasmodesmata, these regions were also deficient in osmiophilic material except for a faint signal perpendicular to some pores traversing the wall (Fig. 3B). Despite these alterations, the centrifugal orientation of chloroplasts was retained in double mutants, and there were no obvious changes in plastid abundance or morphology (Fig. 3C). Interestingly, a similar phenotype was observed in the hypodermis but not the endodermis of double mutant roots, which were largely indistinguishable from WT (Supplemental Figure S11).

**Figure 3.**
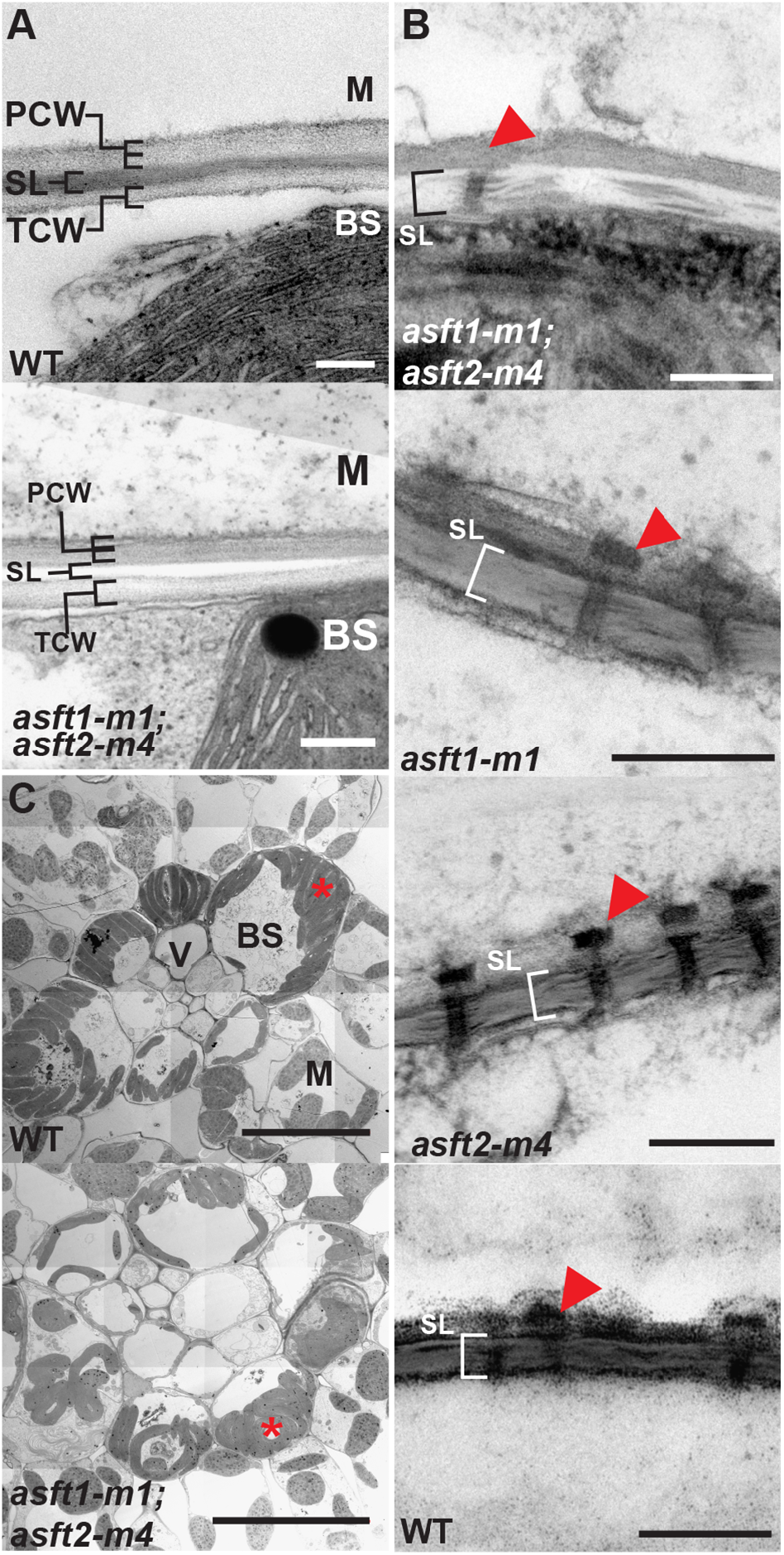
Bundle sheath cell wall ultrastructure is compromised in double mutants. **A.** Transmission electron micrographs of the bundle sheath/mesophyll (BS/M) interface of wild type (WT) and double mutant (*asft1-m1; asft2-m4*). Suberin lamellae (SL) are present between the primary cell wall (PCW) and tertiary cell wall (TCW) in double mutants but are devoid of electron-opaque material and irregular in thickness. 16,000x magnification. Scale bars denote 200 nm. **B.** Polylamellate SL adjacent to plasmodesmata. Arrowheads denote mesophyll-specific caps on plasmodesmata at the BS/M interface. The structure is present in double mutants but occluded in the image presented. Magnifications are 25,000x (*asft1-m1; asft2-m4* and WT), 40,000x (*asft1-m1*), and 31,500x (*asft2-m4*). Scale bars denote 200 nm. **C.** Overview of minor vein anatomy. Representative cells of the BS, M, and vasculature (V) are labeled. Asterisks denote centrifugally positioned chloroplasts within the BS. Magnifications are 800x for WT and 1600x for double mutant. Scale bars denote 20 μm.

As we did not observe a significant increase in aliphatic monomer deposition in double mutant BS strands (Figure 2 and Supplemental Figures S6–S8). we investigated whether the electron lucent regions could be comprised primarily of negative space produced by tearing of the cell wall in the vicinity of the SL. Chromium trioxide was utilized as an alternative fixative, as this reagent successfully discriminated between aliphatic material and negative space in bacterial cell walls (Berg, 1994). WT leaf samples fixed in chromium trioxide had SL with typical “tramline” ultrastructure comparable to samples fixed with osmium tetroxide (Fig. 4A). Conversely, chromium trioxide-fixed leaf sections of the double mutant exhibited significant tearing of both the primary and tertiary cell walls immediately adjacent to the SL (Fig. 4, B and C). Interestingly, the entire SL separated from the polysaccharide cell walls as a discrete unit, and the internal electron lucent region of the lamella was largely indistinguishable from WT (Fig. 4, B and C). In radial cell walls of adjacent bundle sheath cells, SL exhibited a normal “tramline”-architecture (Fig. 4D). Thus, the double mutant specifically compromises the interfaces between the SL and adjoining polysaccharide cell walls, and the *ZmAsft* genes have little effect on the internal patterning of the SL.

**Figure 4.**
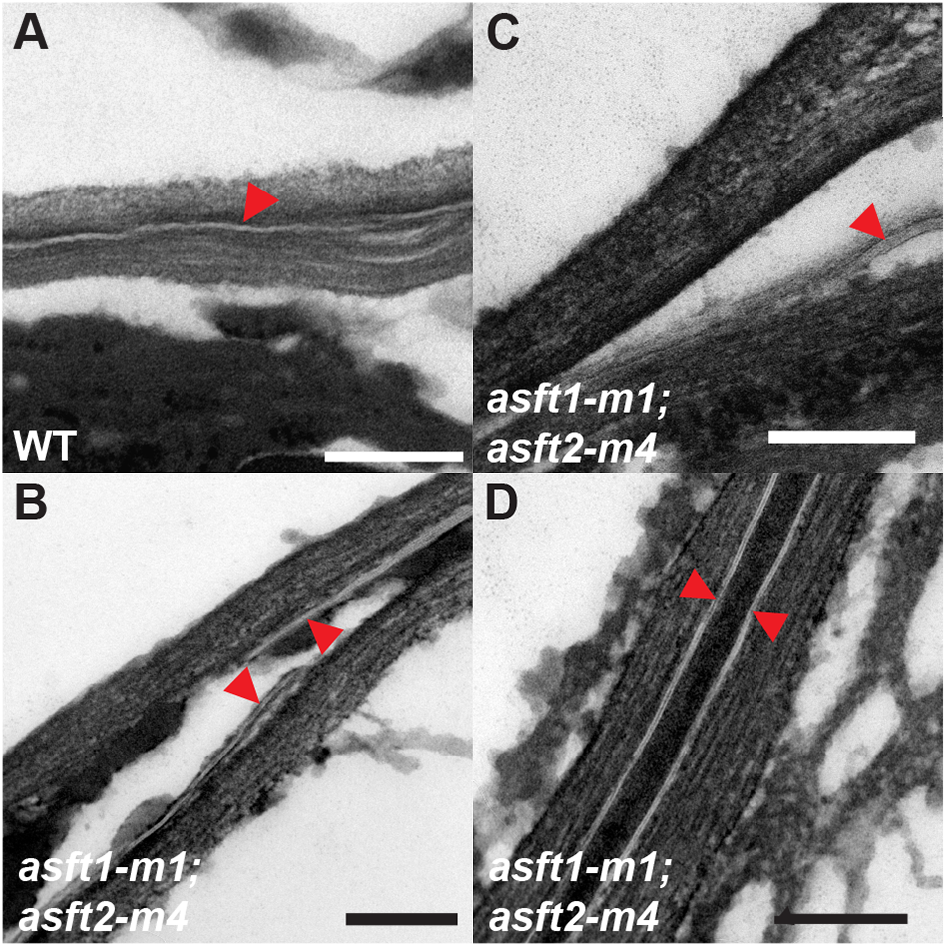
The ultrastructural defect in double mutants is localized to the suberin lamella-polysaccharide cell wall interface. **A.** Transmission electron micrographs of WT BS/M interface. Arrowheads denote SL. 16,000x magnification. **B.** Double mutant BS/M interface. The SL has torn away from the adjoining cell walls as a discrete band. 25,000x magnification. **C..** Double mutant BS/M interface near a cell corner. 31,500x magnification. **D.** Radial cell walls of two adjacent BS cells in the double mutant. The SL are parallel “tramlines” with no evidence of distortion or shearing. 31,500x magnification. All scale bars are 200 nm.

### The ultrastructure and barrier properties of the double mutant cuticle are indistinguishable from wild type

In maize leaves, the suberized BS cells of minor veins are separated from the epidermal cuticle by a single layer of mesophyll cells. As the cuticle is the primary apoplastic barrier to non-stomatal water loss (reviewed in Yeats and Rose 2013), it would be difficult to attribute a change in leaf gas exchange to the ultrastructural defect in the BS if the cuticle was also compromised. Thus, we compared the ultrastructure and barrier properties of the leaf cuticle between WT and *asft* mutants. In contrast to the altered BS ultrastructure, both the cuticle proper and the adjoining polysaccharide cell wall were indistinguishable between mutants and WT (Fig. 5A).

**Figure 5.**
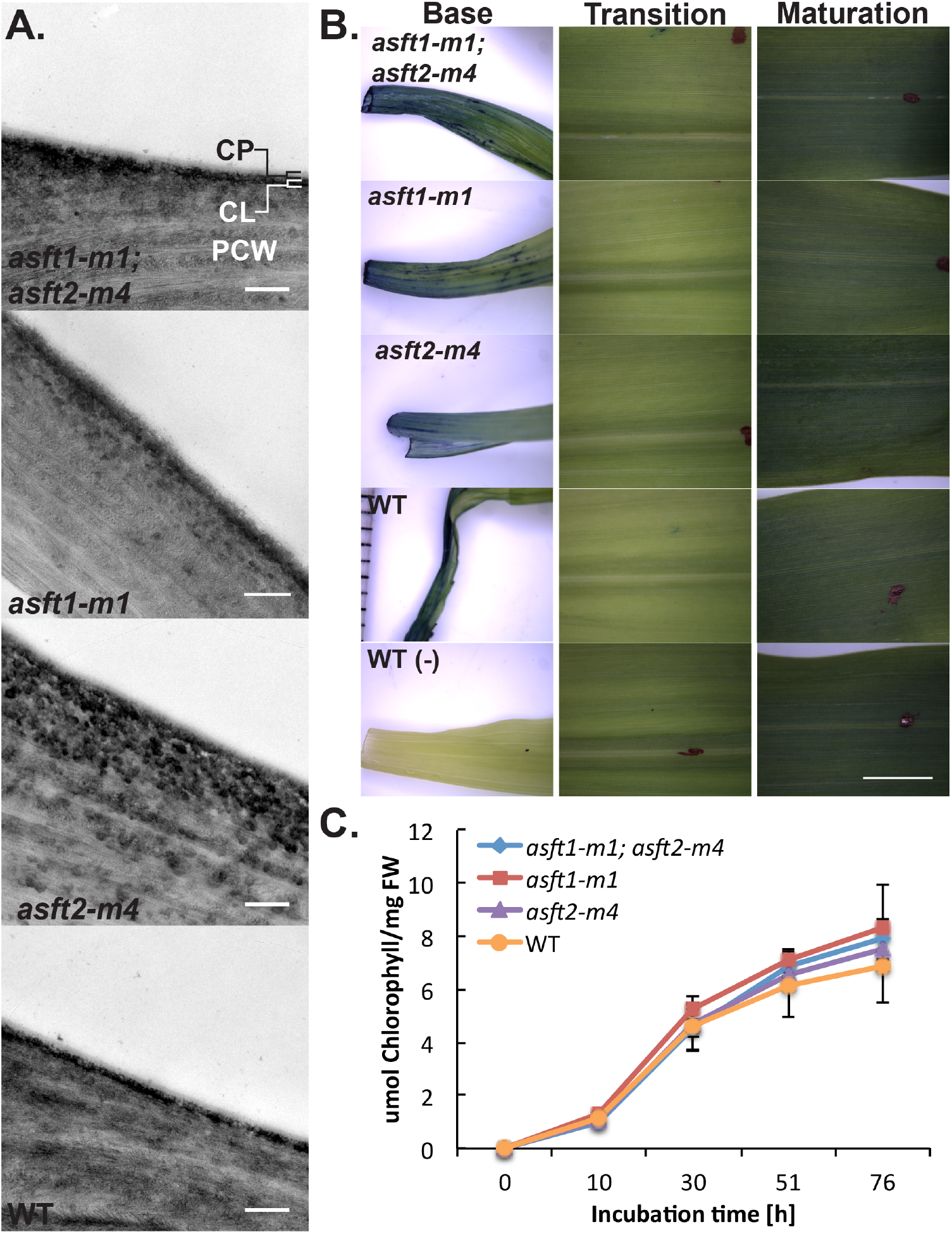
The ultrastructure and barrier properties of the double mutant leaf cuticle are indistinguishable from wild type. **A.** Transmission electron micrographs of the epidermal outer tangential wall in WT and *Zmasft* mutants. The ultrastructure of the cuticle proper (CP), cuticular layer (CL), and polysaccharide primary cell wall (PCW) are indistinguishable between genotypes. Magnifications are 16,000x except for *asft1-m1* (20,000x). Scale bars denote 200 nm. **B.** Toluidine Blue O stains the leaf laminae near the Base in all genotypes, but is excluded from the Transition and Maturation zones. Red ink dots in Transition and Maturation panels denote the point of emergence and the midpoint of the exposed lamina, respectively. Scale bar denotes 50 mm. **C.** Chlorophyll leaching kinetics are indistinguishable between genotypes. Data are presented as average μmoles of chlorophyll per mg fresh weight with standard deviations for 10 biological replicates per genotype.

We utilized Toluidine Blue staining and chlorophyll leaching assays to assess barrier function. In both WT and mutant leaves, Toluidine Blue staining was limited to basal portions of 10-day old third leaves adjacent to the excision site, with no staining observed in the transition zone or in mature source tissue midway between the point of emergence and the leaf tip (Fig. 5B). Likewise, the chlorophyll leaching kinetics for whole third leaf laminae were indistinguishable between genotypes (Fig. 5C; one-way ANOVA, all *p* > 0.1). Thus, the *ZmAsft* genes are redundantly essential for normal BS cell wall ultrastructure but dispensable for normal cuticular ultrastructure and barrier function.

### The double mutant has no effect on CO_2_ concentration or plant growth under hypoxia

To evaluate whether the chemical and ultrastructural defects described above compromised the C_4_ CCM, we measured CO_2_ response curves and measured online ^13^C isotope discrimination (Δ^13^C). Double mutants and WT plants did not differ in the magnitude of the initial slope of the *A-C*_i_ curves (Fig. 8A). There was also no difference in net CO_2_ assimilation (*A*_net_) at any measured internal CO_2_ concentration (*C*_i_) in either the CO_2_-limited phase or in the Rubisco-limited phase of the *A-C*_i_ curve (Fig. 8A). Overall, the double mutant did not affect *A*_net_ at any ambient CO_2_ partial pressure. However, at high ambient CO_2_ concentration (*C*_a_), *C*_i_ (and *C*_i_/*C*_a_) were lower because stomatal conductance (*g*_s_) decreased (Fig. 8B). Likewise, on-line Δ^13^C was not different between genotypes (Fig. 6B).

**Figure 6.**
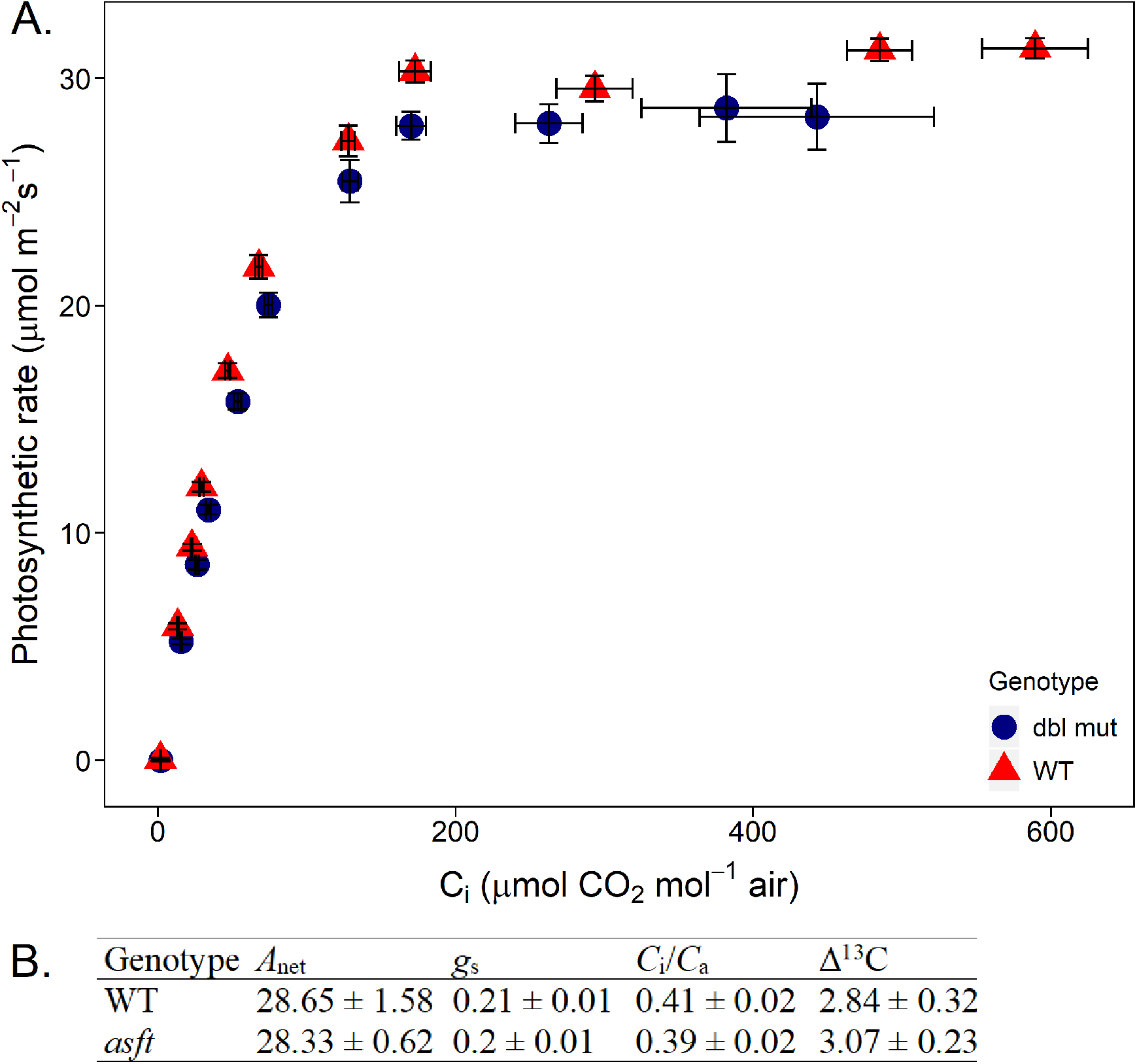
A-Ci curves (A) and on-line photosynthetic ^13^C discrimination (Δ^13^C) (B). **A.** A-Ci curve of net CO_2_ assimilation rate (A_net_) versus intracellular CO_2_ concentration (C_i_). Net CO_2_ assimilation does not differ between wild type (WT, red) and *asft1-m1, asft2-m4* double mutants (dbl mut, blue) at any measured CO_2_ concentration. There is no significant difference in isotopic discrimination or intercellular CO_2_ concentration between double mutant and WT.

The absence of a measurable photosynthetic gas exchange phenotype could indicate either insufficient damage barrier function or that the barrier is compromised but SL are dispensable for the C_4_ CCM. Thus, we performed a waterlogging treatment to evaluate whether *ZmAsft* function is required to maintain growth during hypoxia, where the suberized exodermis constitutes the primary apoplastic barrier against radial oxygen loss (Abiko *et al.*, 2012). Although we observed a significant treatment effect for all growth parameters (two-way ANOVA; all *p* < 0.01), there was no genotype effect within treatment groups for any measured parameter except for leaf width in control plants (Supplemental Table S2). Thus, the ultrastructural defect in double mutant roots does not substantially attenuate plant growth relative to wild type under hypoxic growth conditions. This implies that mutating the *ZmAsft* genes does not cause sufficient damage to the SL to evaluate whether these structures regulate gas exchange in either roots or leaves.

### Water flux and stomatal conductance are enhanced in *Zmasft* double mutants

Pressure-volume curves were measured to evaluate the effect of the *Zmasft* double mutant on leaf water relations from plants grown under both high (1200 μmol quanta/m^2^/s) and low (300 μmol quanta/m^2^/s) light conditions. Double mutants had significantly lower relative water content at the turgor loss point (RWC_TLC_; 0.871 ± 0.01) than WT (0.916 ± 0.005), but light intensity had no effect (2-way ANOVA PAR x genotype *p* < 0.001; Fig. 7A), meaning that the double mutant lost more water in reaching TLP. In contrast, leaf water potential at the turgor lost point (*ψ*_TLP_) did not differ between genotypes (*P* = 0.08). The relative leaf capacitance at full turgor (*c*_ft_) was also elevated in the double mutant (0.16 ± 0.014) compared to WT (0.097 ± 0.008; 2-way ANOVA PAR x genotype *p* < 0.05; Fig. 7C). Under high light conditions, the bulk modulus of elasticity (ε), which bulk modulus estimates the average cell wall rigidity across the leaf (Koide *et al.*, 2000), was significantly lower in double mutants (6.32 ± 0.80) than WT (11.61 ± 0.72; 2-way ANOVA PAR x genotype *p* < 0.0001; Fig. 7B). Saturated water content (SWC) was slightly higher in the double mutant leaves (7.64 ± 0.28) relative to WT (7.06 ± 0.29; 2-way ANOVA PAR x genotype *p* = 0.047). A gas exchange experiment showed that both transpiration rate (*E*) and stomatal conductance per unit stomata (*g*_s_/stomata) were higher in double mutants than WT at high light conditions under low and high relative humidity (3-way ANOVA PAR x RH x Genotype, *p* < 0.0001; Fig. 8, A and B).

**Figure 7.**
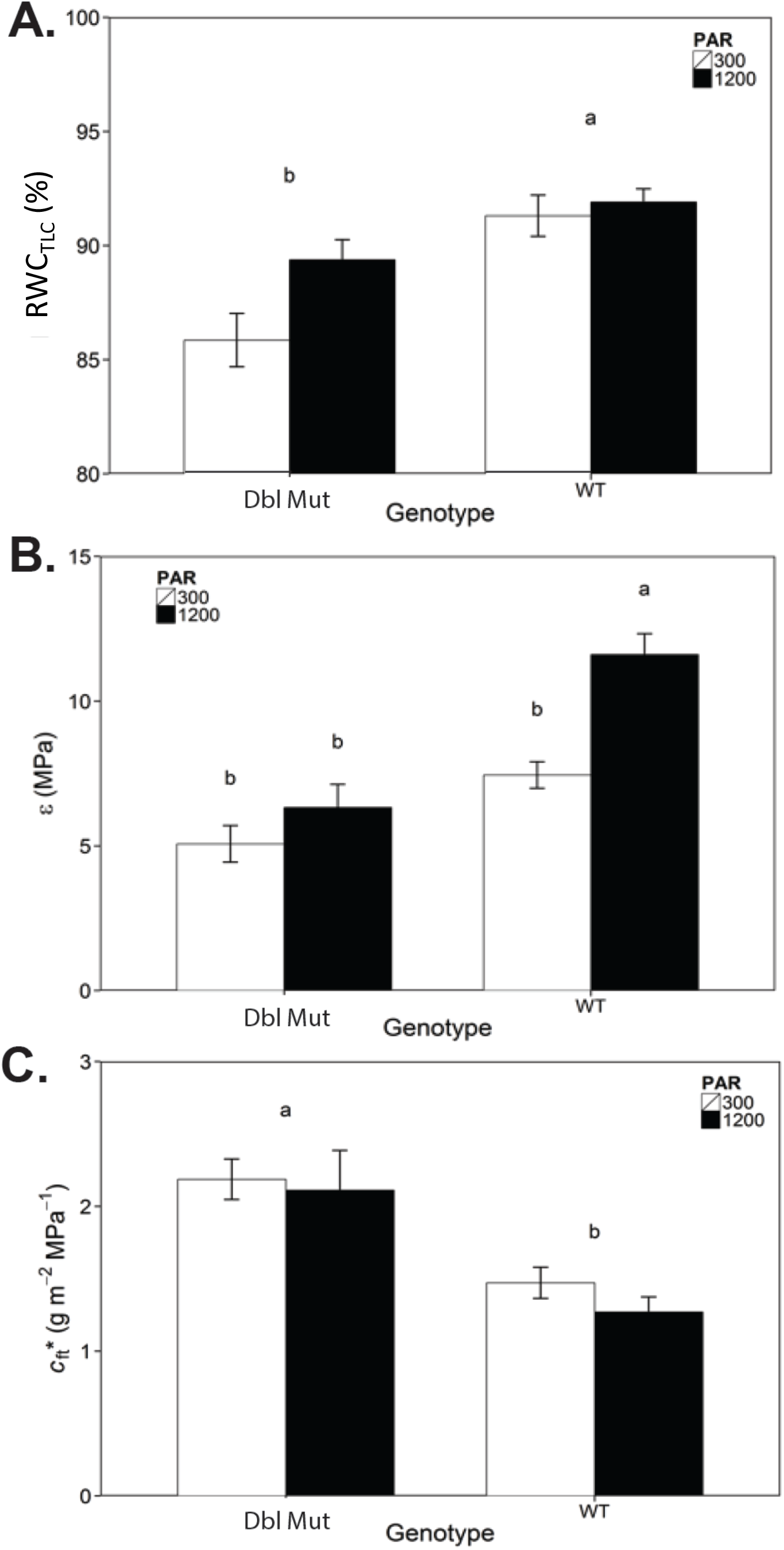
Relative **c**apacitance was elevated and modulus of cell wall elasticity (lower elasticity) was low in double mutants. **A.** Relative water content at the leaf turgor loss point (RWC_TLP_) of *asf1-m1; asft2-m4* double mutants (Dbl Mut) and isogenic wild types (WT) cultivated under high irradiance (1200 μmol photons/m^2^/s) or low irradiance (300 μmol photons/m^2^/s). For all panels, different letters indicate a significant genotype effect. **B.** Bulk elastic modulus (ε) of WT and double mutant. Significantly more water is lost per *ψ*_p_ in the double mutants grown under high irradiance relative to double mutants. There is no genotype effect under low irradiance. **C.** Absolute capacitance at full turgor (c_ft_) is significantly higher in double mutants relative to WT under both irradiance regimes.

**Figure 8.**
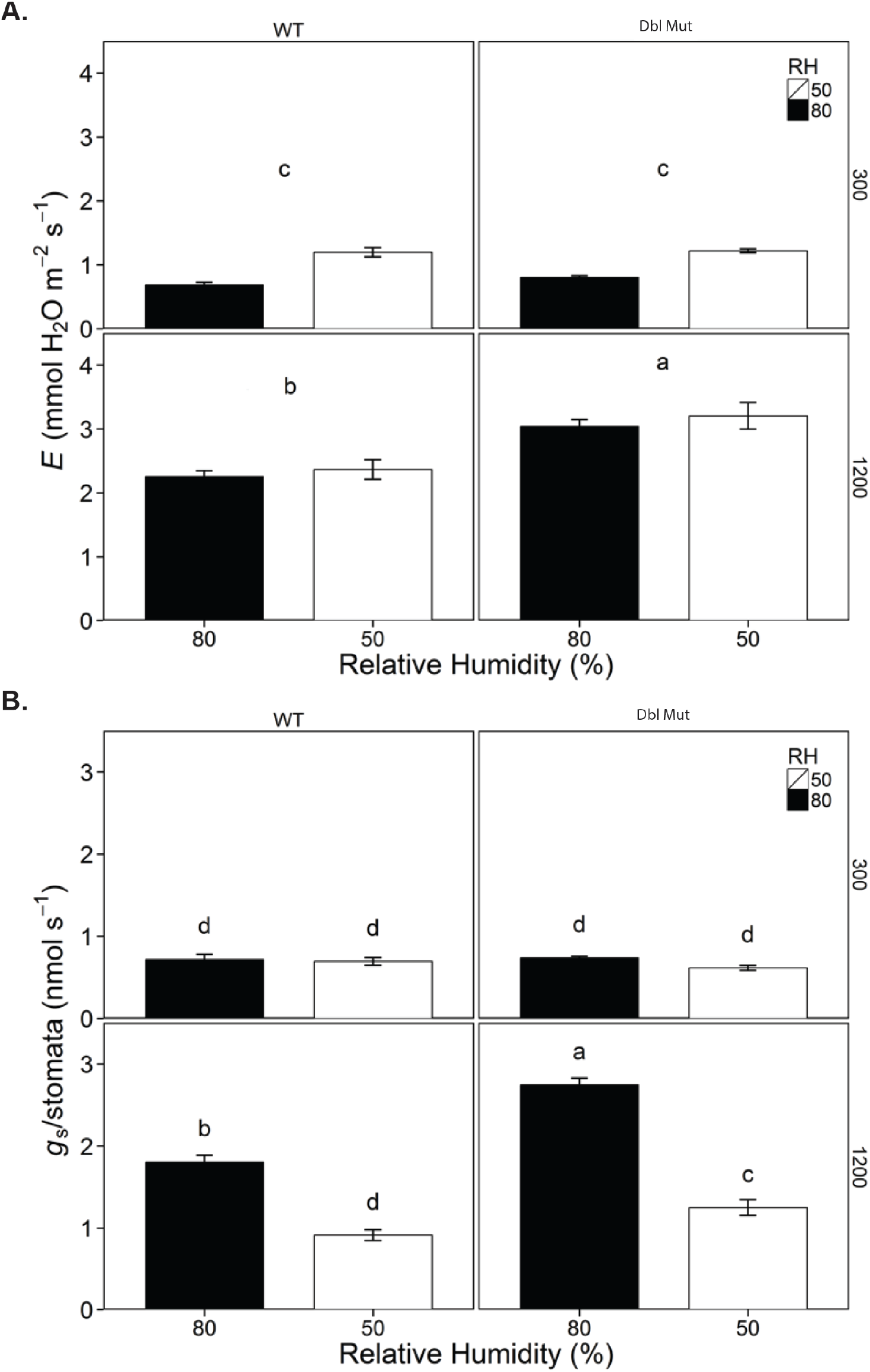
Transpiration rate and stomatal conductance were elevated in double mutants. **A.** Under high irradiance, transpiration flux (E) is elevated in double mutants relative to wild type at both 50% and 80% relative humidity. There is no genotype effect for either humidity regime at low irradiance. **B.** Stomatal conductance normalized by stomatal number (gs/stomata) is elevated in double mutants relative to WT under high irradiance at both 50% and 80% relative humidity. There is no genotype effect for either humidity regime at low irradiance.

## DISCUSSION

### *Zmasft* double mutants are deficient in aliphatic suberin monomers

In order to functionally dissect bundle sheath suberization in an NADP-ME-type C_4_ grass, we identified two paralogously duplicated maize homologues of Arabidopsis *ALIPHATIC SUBERIN FERULOYL TRANSFERASE, ZmAsft1* and *ZmAsft2*, and generated an allelic series of double mutants using *Dissociation* transposons. *asft1-m1*; *asft2-m4* double mutant polyesters were strongly and specifically deficient in C_22:0_-C_30:0_ ω-OH VLCFA relative to both single mutants and WT in leaves and roots (Figure 2; Supplemental Figures S6 and S7). Purified cuticle samples could not be recovered for analysis, and so a pleiotropic leaf cutin defect cannot be ruled out. However, maize suberin is enriched in VLCFA relative to cutin (Espelie and Kolattukudy, 1979), and ω-OH VLCFA are strongly attenuated in purified BS samples (Supplemental Figure S8). This aliphatic suberin deficiency was qualitatively similar to *Atasft* mutant seed coat polyesters, which exhibited a slight reduction in C_22:0_-C_24:0_ ω-OH VLCFA, the longest chain ω-OH FA present in Arabidopsis suberin (Molina *et al.*, 2010). Conversely, *StFHT*-RNAi lines were strongly deficient in C_18:1_ ω-OH LCFA (Serra *et al.*, 2010), potentially reflecting variation in substrate specificity between ASFT proteins. Esterification of ferulic acid to 26- and 28-carbon ω-OH VLCFA was also reported in waxes of oat caryopses, although the causal gene has not been identified (Daniels and Martin, 1968).

Total esterified ferulic acid in mutant Arabidopsis and potato suberin was reduced by 50-90% (Gou *et al.*, 2009; Molina *et al.*, 2009; Serra *et al.*, 2010). Although the aliphatic defect of the *Zmasft* double mutant was consistent with model dicots, the absence of any stoichiometric reduction in HCA was unexpected (Figure 2; Supplemental Figures S6–S8). Both *p*-Coumaric and ferulic acid esters of ω-OH FA have been identified by partial depolymerization of potato periderm suberin and tomato (*Solanum lycopersicum*) fruit cutin (Graça and Santos, 2007; Graça and Lamosa, 2011), and in maize leaves, both *p*-coumaric and ferulic acids were sufficiently abundant to couple to the deficient ω-OH VLCFA in a 1:1 stoichiometry (Molina et al., 2009). However, the fraction of HCA specifically associated with suberin rather than GAX or lignin within the type II cell wall of maize was unknown at the outset of this study. Dilute acid fractionation of the cell wall indicated that a very small fraction of the total ester-linked *p*-coumarate in our samples was associated with GAX (8.8%; Supplemental Figure S9D). Unlike ferulic acid, *p*-coumarate is extensively esterified to maize lignin as a pendant side-chain, and so the majority of *p*-coumarate is likely associated with lignin (Mueller-Harvey et al., 1986; Ralph et al., 1994). Conversely, 86-90% of total cell wall ferulate in developing leaves is associated with GAX rather than suberin (Supplemental Figure S9C). The estimated quantity of suberin-associated ferulate, 10-14% (1.7-2.3 nmol/mg DW) is in good agreement with the reduction in ω-OH VLCFA (1.5 nmol/mg DW) in double mutants. Interestingly, mutants of Arabidopsis *CYP86B1*, which is required for ω-OH VLCFA biosynthesis, are deficient in ferulic acid, but the orthologous rice *cyp86b3* mutant exhibits no reduction in ferulate despite a complete absence of ω-OH VLCFA in root suberin (Waßmann, 2014). Although a direct covalent linkage between

HCA and ω-OH VLCFA remains to be demonstrated for grass suberin, these results indicate that changes in suberin-associated HCA content may be challenging to quantify in grass mutants due to the large proportion of these linkages associated with the Type-II polysaccharide cell wall.

### The ultrastructural defect of *Zmasft* double mutants differs from model dicots

Models of the suberin polyester proposed prior to the characterization of Arabidopsis *ASFT* frequently included ferulic acid as an essential structural component of the electron-opaque lamellar bands and the interface with the polysaccharide cell wall (Kolattukudy, 1980; Bernards, 2002; Graça and Santos, 2007). However, both lamellar ultrastructure and cell wall cohesion were unaffected in Arabidopsis roots and potato tuber periderm of *Atasft* and *Stfht* mutants (Molina *et al.*, 2009; Serra *et al.*, 2010). Conversely, the BS cell wall ultrastructure of the *Zmasft* double mutant is consistent with earlier structural models. BS cells rupture during mechanical shearing, and the SL tear away from adjoining polysaccharides as discrete units during preparation for TEM (Figures 3 and 4; Supplemental Figure S10). Although double mutants exhibited a significant reduction in ω-OH VLCFAs without a detectable stoichiometric reduction in HCAs (Figure 2; Supplemental Figures S6 and S8), it is likely that an aromatic monomer deficiency underlies the ultrastructural defect. Separation of the wall into discrete layers at interfaces between cellulose microfibrils and electron-translucent suberin lamellae was also reported for secondary walls of green cotton (*Gossypium hirsutum*) fibers treated with aminoindan-2-phosphonic acid (AIP), an inhibitor of phenylalanine ammonia lyase (Fig. 4 of Schmutz *et al.*, 1993). The authors attributed the ultrastructural defect to a deficiency in caffeic acid, which was esterified to the ω-OH FA side chains of acyl-glycerol oligomers (Schmutz *et al.*, 1993). The predominant aliphatic monomer of green cotton suberin is 22-OH docosanoic acid; thus, VLCFA-HCA oligomers are a crucial structural component of the suberin polyester in this cultivar (Ryser *et al.*, 1983; Yatsu *et al.*, 1983). Substantial accumulation of caffeic acid esters in either maize suberin or suberin-associated waxes has not been reported to date (Supplemental Figures S7 & S8; Zeier et al., 1999; Kosma et al., 2015). Of the two predominant HCAs identified in this study, acyl *p*-coumarate esters are less likely than acyl ferulates to contribute directly to a cell wall adhesion defect. In Type II cell walls, ferulate esters of GAX, but not p-coumarate esters of lignin, readily dimerize *in muro* and are major contributors to cell wall adhesion and biomass recalcitrance (Grabber *et al.*, 1998 & 2000; Hatfield *et al.*, 2008). Thus, we hypothesize that ZmAsft1 and ZmAsft2 are ω-OH VLCFA:feruloyl-CoA acyltransferases required for cohesion between SL and feruloylated cell wall polysaccharides.

In contrast to the BS ultrastructural defect, the root endodermal and hypodermal layers unexpectedly showed little to no change in cell wall cohesion, consistent with the phenotypes of previously characterized *asft* mutants (Supplemental Figure S11). In AIP-treated green cotton fibers, the secondary cell walls were comprised predominately of cellulose (95% by dry weight) plus a very small proportion (~2.6%) of HCAs and lignin-like polymers (Timpa and Triplett, 1992; Fan *et al.*, 2009). The minor vein BS cells from several grasses, including maize, fluoresce green rather than blue in the presence of 100 mM ammonium hydroxide (pH 10.3), suggesting that this tissue also contains HCAs but little lignin (Harris and Hartley, 1976). Conversely, the root endo- and exodermis, as well as major vein BS cells, fluoresced blue and reacted positively with phloroglucinol-HCl and Maulë Reagent (data not shown). Thus, lignification of adjoining cell walls may stabilize the double mutant SL in roots, although it remains to be determined whether this involves a direct covalent interaction or a general reinforcement of the entire wall against mechanical disruption. BS cells surrounding major veins were not analyzed in this study, and it will be interesting to determine whether the cell wall cohesion defect extends to these cells.

### *Zmasft* double mutants do not compromise suberization sufficiently to assess its role in gas exchange

*ZmAsft1* and *ZmAsft2* are not necessary for CO_2_ assimilation and do not affect the C_4_ CCM. In C_4_ plants, Rubisco is sequestered within the BS cells, and PEPC carries out the initial carboxylation reaction required for CO_2_ capture (reviewed in Sage, 2004). PEPC exhibits a lower degree of ^13^C isotope discrimination relative to Rubisco, leading to a reduced rate of ^13^C isotope discrimination in C_4_ plants relative to C_3_ species (Farquhar *et al.*, 1989). Increased bundle sheath permeability would theoretically permit direct diffusion of atmospheric CO_2_ to Rubisco, increasing fractionation of the heavy isotopologue ^13^CO_2_ relative to ^12^CO_2_ (elevated ^13^CO_2_ isotope discrimination %) while reducing the ratio of intercellular to ambient CO_2_ concentration due to back-diffusion of captured CO_2_ to the mesophyll (C_i_/C_a_; Henderson *et al.*, 1992). The fact that the mutation did not influence *A*_net_ or ^13^C isotope discrimination indicates either that suberin is not an important factor in CO_2_ movement in the leaf, or that the disruption to the SL ultrastructure in the Zmasft double mutant does not compromise the integrity of the apoplastic diffusion barrier.

The absence of a measurable growth difference between double mutants and wild-type siblings under waterlogging stress supports the latter hypothesis. Molecular genetic data from the first suberin biosynthesis mutants characterized in rice provide compelling evidence that SL and aliphatic suberin, in particular, are required to form diffusion barriers to O_2_, at least in roots. The *reduced culm number1 (rcn1/Osabcg5)* mutant was recently demonstrated to encode an ABCG half transporter essential for elongation of rice roots in stagnant, deoxygenated medium (Yasuno *et al.*, 2009; Shiono *et al.* 2014a). *RCN1/OsABCG5* expression is strongly and specifically induced by hypoxia treatment in the endo- and hypodermal layers of nodal roots, and total aliphatic suberin content is halved in the mutant (Shiono *et al.*, 2014a). Intriguingly, neither enhanced lignification of the subtending sclerenchyma proximal to the cortex nor elevated HCA deposition was sufficient to prevent the penetration of the apoplastic tracers periodic acid and berberine hemisulfate into the cortex (Shiono *et al.*, 2014a). Although diffusional permeability of O_2_ was not measured directly, these data suggest a specific role for aliphatic suberin in the formation of diffusion barriers in roots. The recent characterization of the rice VLCFA ω-hydroxylase CYP86B3 supports this conclusion. Roots of *Oscyp86b3* are entirely deficient in ω-OH VLCFA derivatives, and establishment of a tight barrier to radial O_2_ loss is significantly delayed relative to WT in roots cultivated in well-aerated medium (Waßmann, 2014). An uncharacterized compensatory mechanism reverses this effect under hypoxia and is there is no genotype effect on biomass accumulation after an 18d stagnant medium treatment (Waßmann, 2014). The ω-OH VLCFA defect and normal biomass accumulation of *Oscyp86b3* under waterlogging stress are highly reminiscent of the *Zmasft* double mutant. Biomass differences after a period of waterlogging stress correlate with differences in radial O_2_ loss between *Hordeum* and *Zea* species with and without strong constitutive exodermal gas exchange barriers (Malik *et al.*, 2011; Abiko *et al.*, 2012). Taken together, these data suggest that the significant attenuation of ω-OH VLCFA in these mutants is not sufficient to abolish barrier function, and that stronger mutants are necessary to functionally dissect suberization. In particular, the *rcn1/Osagcg5* mutant suggests that interfering with export of aliphatic suberin to the apoplast may be a viable approach, though it remains to be determined whether the pleiotropic morphological phenotypes of this mutant are caused directly by its suberization defect (Shiono et al., 2014a).

### The *Zmasft* double mutant has a subtle effect on leaf water movement

Altered apoplastic water permeability is one of the most prevalent phenotypes reported for suberin mutants. Disruptions to VLCFA elongation by β-ketoacyl-CoA synthases (*StKCS6;* Serra et al., 2009b), LCFA ω-hydroxylation (*AtCYP86A1* and *StCYP86A33;* Hofer *et al.*, 2009; Serra *et al.* 2009a), and suberin feruloylation (*StFHT*; Serra et al., 2010) are all sufficient to increase peridermal water permeance. Root pressure probe experiments with *Atcyp86a1* suggest that both hydrostatic and osmotic hydraulic conductivity are elevated, indicating greater flux through both apoplastic and cell-to-cell pathways (Ranathunge *et al.*, 2011). As reported for epidermal cuticles, suberin-associated wax is the major barrier to peridermal transpiration; extraction of wax from potato tubers increased the water permeance 10-100 fold (Schreiber *et al.*, 2005; Schreiber, 2010). However, the composition of the suberin polyester is also critical, as the *StFHT*-RNAi line exhibited 15-fold higher permeance despite WT levels of total suberin and elevated wax deposition (Serra *et al.*, 2010). The latter study also implies that suberin feruloylation is critical for to restrict uncontrolled diffusion of water, even if the ultrastructure of *StFHT*-RNAi lines appears indistinguishable from WT (Serra *et al.*, 2010).

Leaves of *Zmasft* double mutant are more permissive to water movement, resulting in less control of water loss as observed from greater change in relative water content with leaf water potential (lower ε and RWC_TPL_, higher *c*_FT_). Apoplastic tracer studies indicate that the middle lamellae of radial walls between adjoining bundle and mestome sheath cells form an uninterrupted path for the diffusion of water and solutes out of the leaf vasculature (Evert *et al.*, 1985; Botha and Evert, 1986; Eastman *et al.*, 1988b). In *Zmasft* double mutants, the ultrastructural changes at the interface of the SL and the primary cell wall appear to facilitate apoplastic water movement through this pathway or potentially reduce control of leakage from the vasculature. This greater movement of water out of the vascular bundle provided greater water availability for transpiration. Lower ε means that the double mutant has increased change in water content with *ψ*_p_, and this is interpreted as having higher cell wall elasticity and, in particular, the elastic (reversible) properties of the cell walls. In a solid-state ^13^C NMR study of isolated periderm suberin from *StFHT*-RNAi lines, the cell wall was more stiff due to increased incorporation of recalcitrant polyphenolic material into the cell wall, which caused the cell wall to become more brittle and increased water permeability (Serra *et al.*, 2014). In both *Zmasft* double mutants and *StFHT*-RNAi lines, the mutation resulted in greater cell wall permeability and less restricted apoplastic water movement from the vasculature, but it is unclear if the mechanism is the same. In all, greater water loss to reach TLP, lower RWC_TLP_, and higher *c*_FT_ indicate that the double mutants have less ability to restrict apoplastic water movement. Additionally, this would indicate that the double mutants would be less tolerant of water limitation where greater control over water loss is necessary. As the ultrastructure and barrier properties of the cuticle were not affected by the double mutation (Fig. 5), greater *E* and *g*_s_ likely resulted from greater apoplastic water flow rather than a greater evaporative surface or increased membrane permeability.

## CONCLUSION

Although the suberin biosynthesis pathway is broadly conserved between model dicots and maize, the selection of suitable candidate genes to disrupt barrier function was limited by the availability of *Dissociation* transposons in sufficiently tight linkage to generate targeted insertional mutants at high frequency at the outset of this study. We mutated two functionally redundant paralogues of Arabidopsis *ALIPHATIC SUBERIN FERULOYL TRANSFERASE, ZmAsft1* and *ZmAsft2*, using *Ds* elements and found them to be essential for normal accumulation of omega-hydroxy very long chain fatty acids in bundle sheath suberin. Double mutants exhibited a cohesion defect between bundle sheath suberin lamellae and cell wall polysaccharides indicative of cryptic changes to cell wall architecture. However, the defect was insufficiently strong to characterize the role of bundle sheath suberization in the C_4_ carbon concentrating mechanism. A recently published functional dissection of a rice exodermal suberin mutant suggests that a more severe disruption to the aliphatic suberin polymer is necessary to compromise the gas exchange barrier (Shiono et al., 2014).

The development of monocot-optimized, multiplexed gene editing tools (Cermák et al., 2017) eliminates the restriction of mutagenesis candidates to genes with transposons in linkage to all potential paralogues. Rather than evaluating individual biosynthesis candidates for strong, suberin-specific phenotypes, a more promising approach may be to pursue multiplexed knockouts of suberin transcriptional regulators, which are both distinct from cuticle development (Kosma et al., 2014) and likely to be conserved between model dicots and cereals (Wang et al., 2014). This approach will be the focus of future studies.

## MATERIALS AND METHODS

### Plant Growth Conditions

Unless otherwise indicated, seedlings were germinated in flats (5 columns by 10 rows, 10 cm soil depth) filled with 3:1 (v/v) Metromix 360 (Hummert 10-0356-1):Turface (Hummert 10-2400). At 12-14 days after sowing, seedlings were transplanted into Classic 1000 pots (25.7 cm x 23.2 cm x 20.6 cm) filled with ProMix-BRK 20 potting mix (Hummert 10-2018-1) supplemented with 10 mL fertilizer mix (Osmocote [15-9-12], Tomato Maker, Sprint, FeSO_4_; 1:1: 0.083: 0.06). Beginning 10 days after sowing, plants were watered weekly with a 200 ppm solution of 15-5-15 fertilizer (Calcium-Magnesium LX; Hummert 07-5902-1).

For seedling primary root experiments, kernels were sown in individual Deepot D40L containers (Steuwe and Sons) filled with 1:1 (v/v) Sand (Hummert 10-2210-1):Profile (Hummert 10-2390-1) and cultivated for 10 days in a greenhouse as described below.

Seedlings were germinated in a Conviron BDW growth room with a daily irradiance regime of 550 μmol photons/m^2^/s, a photoperiod of 12 h light:dark cycle, day/night temperatures of 31°C/22°C, and 50% relative humidity. From transplantation to maturity, plants were cultivated in a greenhouse (300 μmol/m^2^/s irradiance, 12h/12h photoperiod, day/night temperature and relative humidity 28 °C/24 °C and 30%/50%) at the Donald Danforth Plant Science Center in Saint Louis, MO.

### Reverse genetic screen using *Ds* transposons

Generation of testcross populations, seedling growth, tissue pooling and deconvolution, and PCR screening were conducted as described in Studer *et al.* (2014) with the following modifications. A total of 450 *Ac*-positive kernels were planted per screen, and the tissue of 15 individual plants was pooled, for a total of 30 pools per screen. All PCR screens were conducted using Platinum Taq HF (Invitrogen) to facilitate long-distance screening of insertions distal to the target gene coding sequence. Individuals containing germinal insertions were backcrossed to T43, and single mutants were cross-pollinated to generate double mutants, which were validated by PCR genotyping (Supplemental Table S3).

### Transcript expression analysis by qRT-PCR

Ten days after sowing, third leaves of 16 ± 1 cm were harvested as described in (Li *et al.*, 2010; Wang *et al.*, 2014). For root experiments, primary roots of 20 cm ± 5 cm were divided into 4 cm increments from the apex and immediately pooled and frozen as described above. Eight plants were pooled per biological replicate, and three to four replicates were collected per genotype.

Total RNA was extracted as in Wang et al., 2014, suspended in RNASecure Resuspension Solution (Ambion; AM7010) and integrity was verified via agarose gel electrophoresis. DNase treatment and cDNA synthesis were conducted as in (Wang et al., 2014) using a 1h reverse transcription reaction at 50°C.

All qPCR reactions were conducted as in Wang et al., 2014 with primer concentrations of 0.4 μM and 4 μL of [1:160] diluted cDNA aliquoted per reaction. Sequences and dilution curve validation of all primers is described in Supplemental Table S3 and Supplemental Table S4. The reference gene primers were previously validated for stable expression in developing leaves (Wang *et al.*, 2014). Threshold cycle values were obtained from the LightCycler software using the Absolute Quantification/2^nd^ derivative maximum method. Normalized relative quantities were calculated using the E^ΔCT^ method as described in Hellemans *et al.* (2007). Statistical significance was determined by a one-way ANOVA followed by a Tukey-Kramer post-hoc test for values giving significant F-statistics (*p* < 0.05).

### Harvesting leaf tissue for cell wall compositional analyses

For all isolated BS strands of 10-day-old plants (Figure 2), 300-450 mg of fresh tissue from fully emerged regions of developing third leaves of 8 individuals were pooled per biological replicate. Leaves were cut into 3 mm square pieces and immediately suspended in 15 mL cold (4°C) grinding buffer 1 (330 mM sorbitol, 300 mM NaCl, 100 mM MgCl_2_, 10 mM EGTA, 10 mM DTT, and 0.5 mM DETC in 200 mM Tris buffer, pH 9.0) and macerated for 30 s using a polytron liquid homogenizer (speed 6), filtered through 60 μm mesh to remove mesophyll protoplasts and cell wall fragments, and re-suspended in 15 mL grinding buffer 2 (350 mM sorbitol, 50 mM EDTA, and 0.1% [v/v] β-mercaptoethanol in 50 mM Tris buffer, pH 8.0). Leaf suspensions were homogenized for 1 minute, filtered, and re-suspended a total of 3 times. Pellets were immediately wrapped in aluminum foil, flash frozen in liquid nitrogen, and stored at −80°C.

For purified bundle sheath strand preparations (Supplemental Figure S9), pellets were prepared as described above and filtered through a stainless steel no. 35 mesh (500 μm) sieve with several washes of MILLI-Q grade water. The eluent, containing isolated bundle sheath strands, was captured by decanting onto a 60 μm mesh. The residual pellet, containing a mixture of cuticle strips and residual bundle sheath strands that did not pass through the sieve, was collected on a separate 60 μm mesh, and both tissues were collected and frozen as described above.

### Leaf Polyester Analysis

Delipidation was conducted according to the Lipid Polyester Analysis protocol of Beisson and colleagues (Li-Beisson *et al.*, 2013), with the following modifications. Following isopropanol boiling, samples were macerated three times for 1 minute each using a PowerGen 500 liquid homogenizer (Fisher) at speed setting 6. All Isopropanol, chloroform:methanol, and methanol washing steps were repeated twice. Samples were stored in a desiccator containing Drierite until a stable dry weight was achieved.

Base-catalyzed transmethylation and silylation were conducted according to (Li-Beisson et al., 2014). 5 μL each of fresh omega-pentadecalactone (10 μg/μL) and methyl heptadecanoate (10 μg/μL) in methanol were injected into each sample immediately prior to transmethylation. Samples were re-suspended in 50 μL chloroform for quantification by GC-FID and identification by GC-MS. Samples (2 μL) were injected onto a TRACE TR-5MS GC column (30 m length x 250 μm ID x 0.25 μm film thickness) using a 1:10 split injection ratio at 320°C (GC-FID) or splitless injection (GC-MS). For GC-FID, the oven was programmed to hold at 140°C for 2 minutes, ramp to 320°C at 5°C/minute, and hold for 10 minutes. Helium carrier gas was supplied at a flow rate of 1.5 mL/min. Monomers were normalized by area ratios using methyl heptadecanoate as an internal standard for fatty acid methyl esters, and omega-pentadecalactone for silylated species as described in (Jenkin and Molina, 2015). For GC-MS, the oven temperature was held at 120°C for 1 minute, then ramped to 320°C at 5°C/minute, followed by a 15 minute hold. Helium carrier gas was supplied at a 1 mL/minute flow rate. Fragmentation spectra were obtained using an Agilent quadrupole detector operating in electron impact mode with an accelerating voltage of 2500, scanning from 35-750 amu at a rate of 0.3 s/scan with a 0.1 s interscan delay and a 5 minute solvent delay at the beginning of each run.

### Electron microscopy

Marginal leaf sections were harvested from the midpoint of fully expanded fourth and fifth leaf laminae of 28d old mutant and WT plants. For roots, 1 cm sections were collected 0.5 cm behind the primary root tip of 10d old seedlings, in the region of first lateral root emergence (8-12 cm from tip), and 2 cm from the root base. Samples were collected in biological triplicate.

Leaf (Figures 3A and 3C) and root samples (Supplemental Figure S11) were hand sectioned into 1 mm fragments in 2 mL of freshly prepared 100mM PIPES buffer (pH 6.8) containing 2% (v/v) glutaraldehyde solution, dehydrated in an ethanol/acetone series, and embedded in Spurr’s resin. For high pressure frozen leaf samples in Figure 3B, and Figure 5, discs were collected from leaf margins using a Harris Uni-Core 1.20, transferred into 100 μm sample planchettes, freeze substituted in 2% osmium tetroxide at (-) 85 C and embedded in Spurr’s resin.

Chromium trioxide preparations were carried out as described in Berg (1994). Leaf tissues were hand sectioned into 1 mm fragments in water, and then transferred into a 10% (v/v) chromium trioxide solution for simultaneous fixation and staining for 1 hour at room temperature. The samples were washed with repeated aliquots of water until the rinse solution became clear, and then dehydrated and embedded as described above. Sample sectioning and TEM were carried out as described in Weissmann et al., 2014.

### Cuticular Permeability Assays

The Toluidine Blue O permeability assay was carried out as described in Tanaka *et al.* (2004) with an extended incubation time for monocot leaves as described in Wu *et al.* (2011). Third leaf laminae of 10d old mutant and WT seedlings were immersed in a 0.05% (w/v) solution of Toluidine Blue O dissolved in distilled water or distilled water (negative control) for 10 minutes, immersed in distilled water to remove residual stain, and blotted dry. Leaves were imaged using a Nikon Dissecting Microscope with bright-field illumination. The experiment was repeated twice with identical results.

The chlorophyll leaching assay was carried out according to Lolle *et al.* (1997). Ten individual third leaves per genotype were weighed and placed into 30 mL of 80% (v/v) ethanol.

Tubes were covered with aluminum foil and incubated on a rocking agitator at room temperature for 3d. At the indicated time-points, absorbance values (A_664_ and A_647_) of a 100 μL aliquot of chlorophyll were measured using a spectrophotometer. Absorbance values were converted to μmoles chlorophyll according to (Lolle *et al.*, 1997) and normalized per unit fresh weight.

### Physiological experiments

Three experiments were conducted to measure (exp. 1) A-Ci curves and on-line ^13^C isotope discrimination, (exp. 2) pressure-volumes curves, and (exp. 3) gas exchange under light intensity and relative humidity treatments. In all three experiments, maize was planted in peat-based potting medium (Sun Gro Horticulture, Sunshine mix LC1) and were grown in Conviron BDW growth chambers. Plants were watered daily and fertilized every 4 days with Peters Pro 20-20-20 initially and later 15-5-15 CalMag.

For the CO_2_ assimilation and on-line ^13^C isotope discrimination experiment (exp. 1), *asft1-m1*; *asft2-m4* double mutants and isogenic wild type were grown under 12 h light:dark, 500 μmol quanta m^2^ s^-2^, 31/21 °C, and ~50 % RH. At four weeks post-sowing, CO_2_ response (*A-C*_i_) curves were measured on the uppermost fully expanded leaf of four plants per genotype as described in Studer *et al.* (2014). On-line ^13^C discrimination experiments were performed on the same plants using an LI6400XT (LI-COR Biosciences) with a LI6400-22 leaf chamber and a LI6400-18 light source coupled to a tunable diode laser absorption spectroscope (TDL-AS, TGA 100A; Campbell Scientific) as described by Ubierna *et al.* (2013).

For the pressure-volume curve experiment (exp. 2), five replicates of each genotype were grown at day and night temperatures of 28°C and 20°C, respectively, and relative humidity was maintained at 70 %. The photoperiod was 16 h with a 1-hour ramp for temperature and light. All plants were grown simultaneously in the same growth chamber with the height of the pots adjusted to maintain the desired light treatments at canopy level from before germination until the conclusion of the experiment. Plants in the high light (HL) group were grown under a photosynthetic photon flux density (PPFD) of 1200 μmol quanta m^-2^ s^-1^ at canopy level, and the low light (LL) group were grown under a PPFD of 300 μmol quanta m^-2^ s^-1^ at canopy level.

Pressure-volume curves were generated using a Scholander pressure bomb as described in Tyree and Hammel (1972). The fifth leaf blade of the maize plants was removed before the lights turned on (predawn) in the growth chamber. Samples were equilibrated for 2 hours at 4 °C in separate plastic bags prior to measurements. Each leaf was weighed, and immediately thereafter each leaf was inserted into the pressure bomb, and the pressure within the pressure bomb was increased slowly until water came out of the cut end of the midvein. The leaf water potential (*ψ*) was the pressure required to force water out of the midvein. This process of weighing and measuring *ψ* was repeated in rapid succession (every 1-3 minutes), while *ψ* decreased substantially between measurements. Once the change in *ψ* between measurements was small, leaf weights and *ψ* were measured at intervals of 4, 10, and 20 minutes for a total of 13 to 17 measurements per leaf.

Relative water content (RWC) was calculated as leaf water volume per dry leaf weight. The inverse leaf water potential (1/*ψ*) was plotted against the water lost from the leaf (1-RWC), and turgor loss point (TLP) was the point of transition between the initial curved portion and the latter linear portion. TLP is the point when the hydrostatic pressure within the cell is equal to atmospheric pressure, so there is no net pressure within the cell wall. Leaf water volume was plotted against *ψ* to calculate saturated water content (SWC) in the initial hydrated portion of the pressure-volume curve including TLP. SWC was the y-intercept of this relationship divided by dry leaf mass. *ψ* is the sum of the water potential from turgor pressure (*ψ*_p_) and osmotic potential (*ψ*_o_); therefore, after the TLP, *ψ_p_* equals zero, so *ψ* equals *ψ*_o_. *ψ*_o_ was calculated for all points by finding the Model II regression of the relationship between 1/*ψ* and (1-RWC) for TLP and the portion after TLP and then using the regression equation to calculate *ψ*_o_ for each (1-RWC) as the negative inverse of [slope * (1-RWC) + intercept]. *ψ*_p_ was the difference between *ψ* and *ψ*_o_. The bulk modulus of elasticity at full turgor (*ε*) determines the rigidity of the cell wall and is the Model II slope when *ψ*_p_ was plotted against RWC for points before and including TLP. Relative capacitance at full turgor (*c*_FT_) was calculated as the Model II slope when RWC was plotted against *ψ* for points before and including TLP.

For the gas exchange experiment (exp. 3), plants were cultivated as described above for the pressure-volume experiment, except that the experiment was replicated in separate growth chambers at two relative humidity levels (50% and 80%). Gas exchange measurements were performed in Pullman, WA, USA, with mean atmospheric pressure of 92.1 kPa. The mid portion of 5^th^ leaf was used for gas exchange measurements. Gas exchange measurements were made on the LI6400XT (LI-COR Biosciences) with a 2×3 cm leaf chamber with a red and blue light emitting diode (LED) light source. Point measurements were made with the growth chamber on the mid-section of the leaf blade, so that point measurements would reflect growing conditions. Leaf chamber conditions were: leaf-to-air vapor pressure deficit of 1 kPa, leaf temperature of 28°C, flow rate of 300 μmol air s^-1^, light intensity at saturating photosynthetic active radiation of 1500 μmol quanta m^-2^ s^-1^, and oxygen partial pressure of 19.3 kPa (21%).

### Gas Exchange Measurements Coupled with Carbon Isotope Discrimination for calculations of mesophyll conductance

For each CO_2_ concentration, 6-8 cycles of TDL measurements were performed. The TDL-AS measured the carbon isotopes in both reference (inlet) and samples (outlet) air from each Licor chamber. The TDL-AS measurements were calibrated for gain and possible offset using two calibration tanks (Liquid Technology), according to Bowling *et al.* (2003) and Ubierna *et al.* (2013). Photosynthetic ^13^C discrimination was calculated as described by Evans *et al.* (1986):

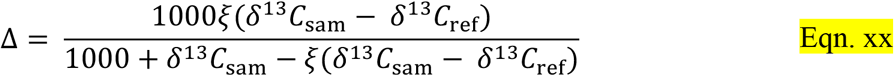

where δ^13^C_sam_ and δ^13^C_ref_ are the carbon isotope composition of the leaf sample (chamber) and reference air of the LI-6400XT. The ξ values were calculated as ξ = *C*_ref_/(*C*_ref_-*C*_sam_), where *C*_ref_ and *C*_sam_ are the CO_2_ concentrations of dry air entering and exiting the leaf chamber, respectively. The delta notation expressed in ‰ was relative to Vienna Pee Dee Belemnite standard.

## List of author contributions

R.A.M. designed the initial research plan under the guidance of T.P.B.; T.N., A.B.C., S.vC., and T.P.B. supervised the experiments; R.A.M., P.E., P.E., S.L.T.,, S.vC., and R.H.B. conducted the experiments and analyzed the data; R.A.M. and P.E. wrote the manuscript and revised it with contributions from all the authors.

Gene identification and mutagenesis, transcript profiling, cell wall analyses, and microscopy were funded by an NSF Plant Genome Research Project Grant (IOS-1127017) to T.P.B and T.N. CO_2_ assimilation and carbon/oxygen isoptope exchange measurements were funded by grants (numbers) issued to A.B.C. and S.vC.

---

**Supplemental Figure S1.**
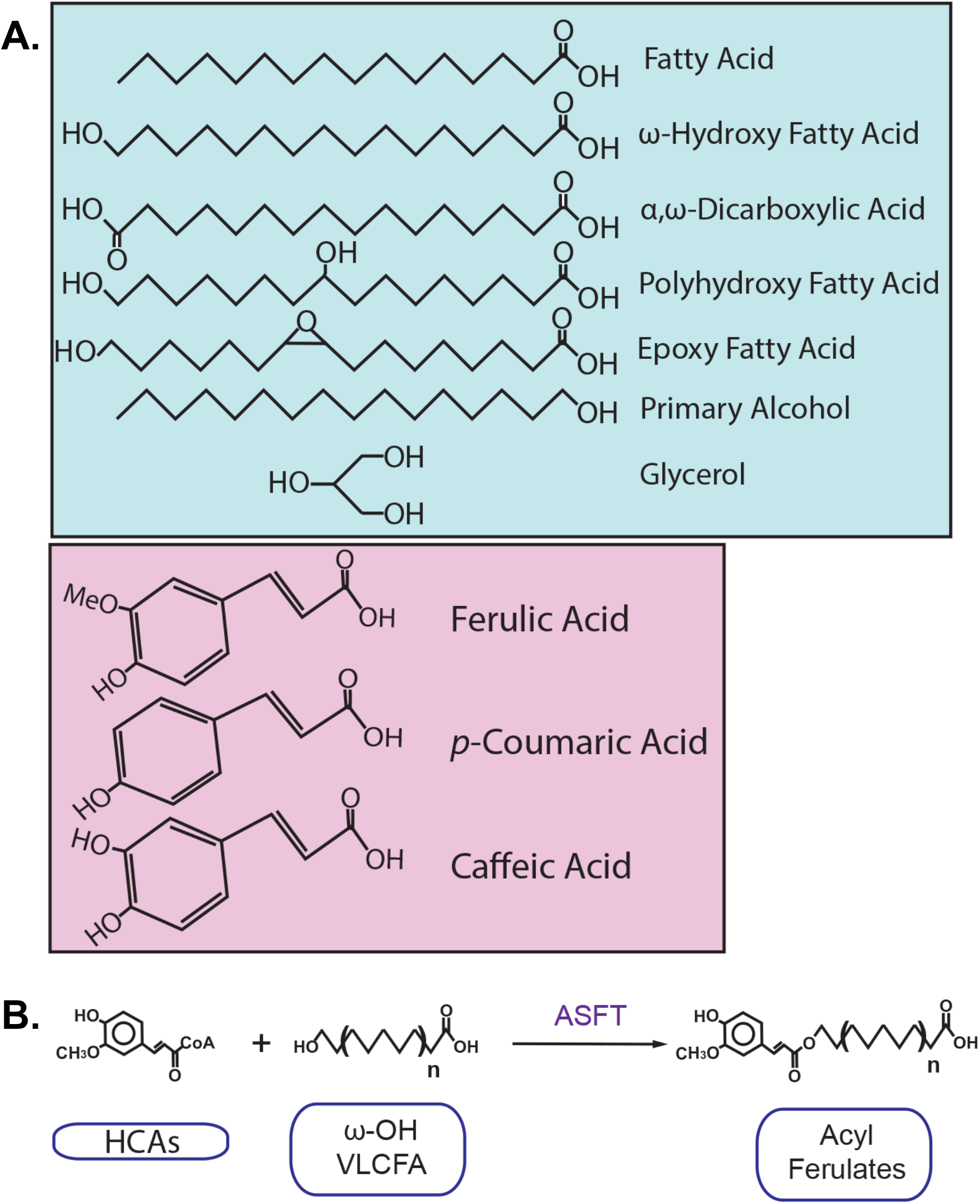
**A.** Structures of representative aliphatic (blue box) and aromatic (purple box) suberin and cutin monomers identified in this study. **B.** Aliphatic Suberin Feruloyl Transerase (ASFT) generates acyl ferulates for suberin polyester synthesis by esterifying hydroxycinnamoyl-CoAs (HCAs) to free hydroxyl groups of omega-hydroxy very long chain fatty acids (w-OH VLCFA) or primary alcohols.

**Supplemental Figure S2.**
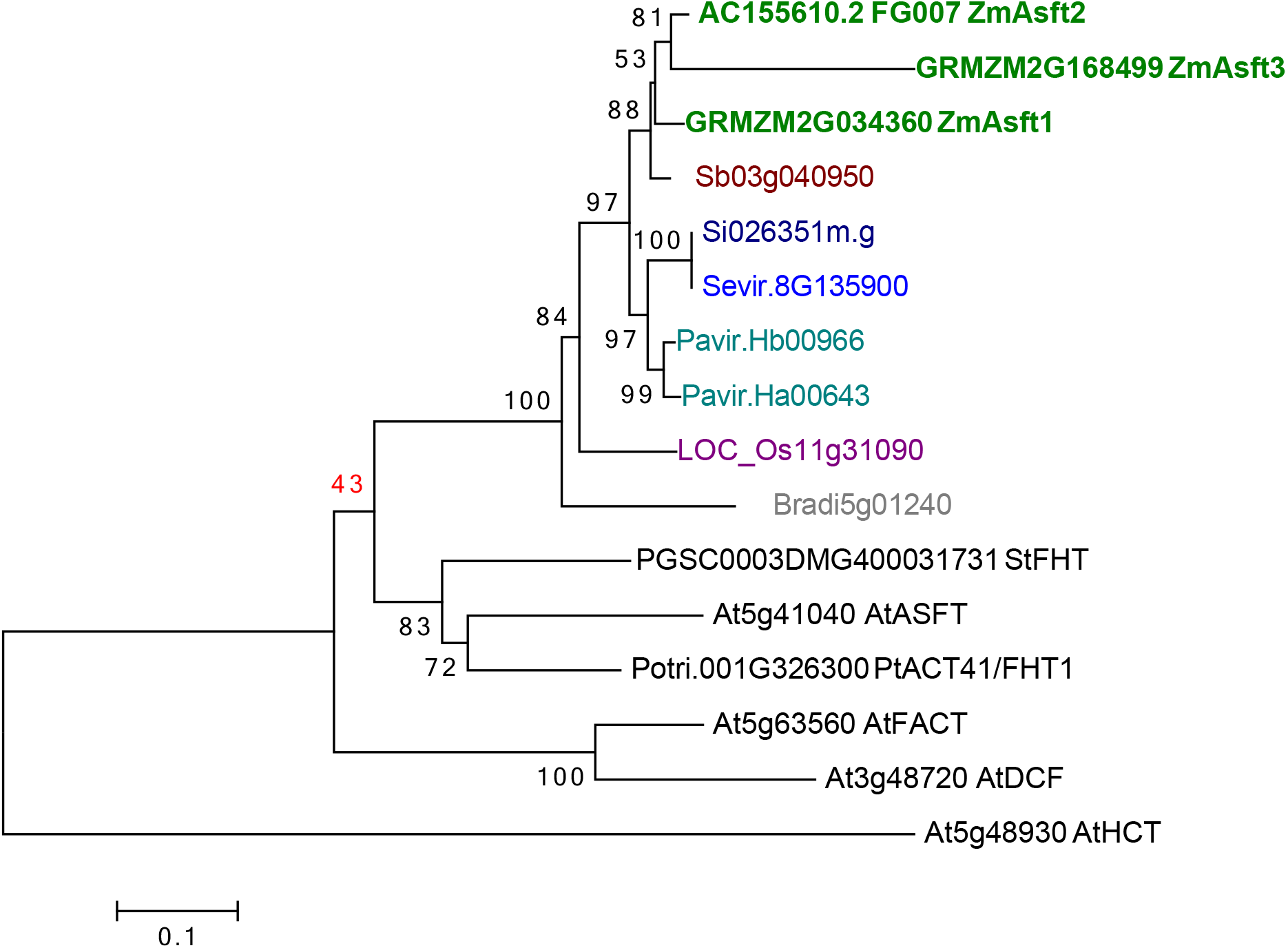
Rooted Neighbor-Joining phylogeny of ASFT sub-clade proteins from grasses and model dicots. AtHCT, a BAHD acyltransferase involved in lignin biosynthesis, serves as the outgroup. Bootstrap support (1000 replicates) is shown next to the corresponding branches.

**Supplemental Figure S3.**
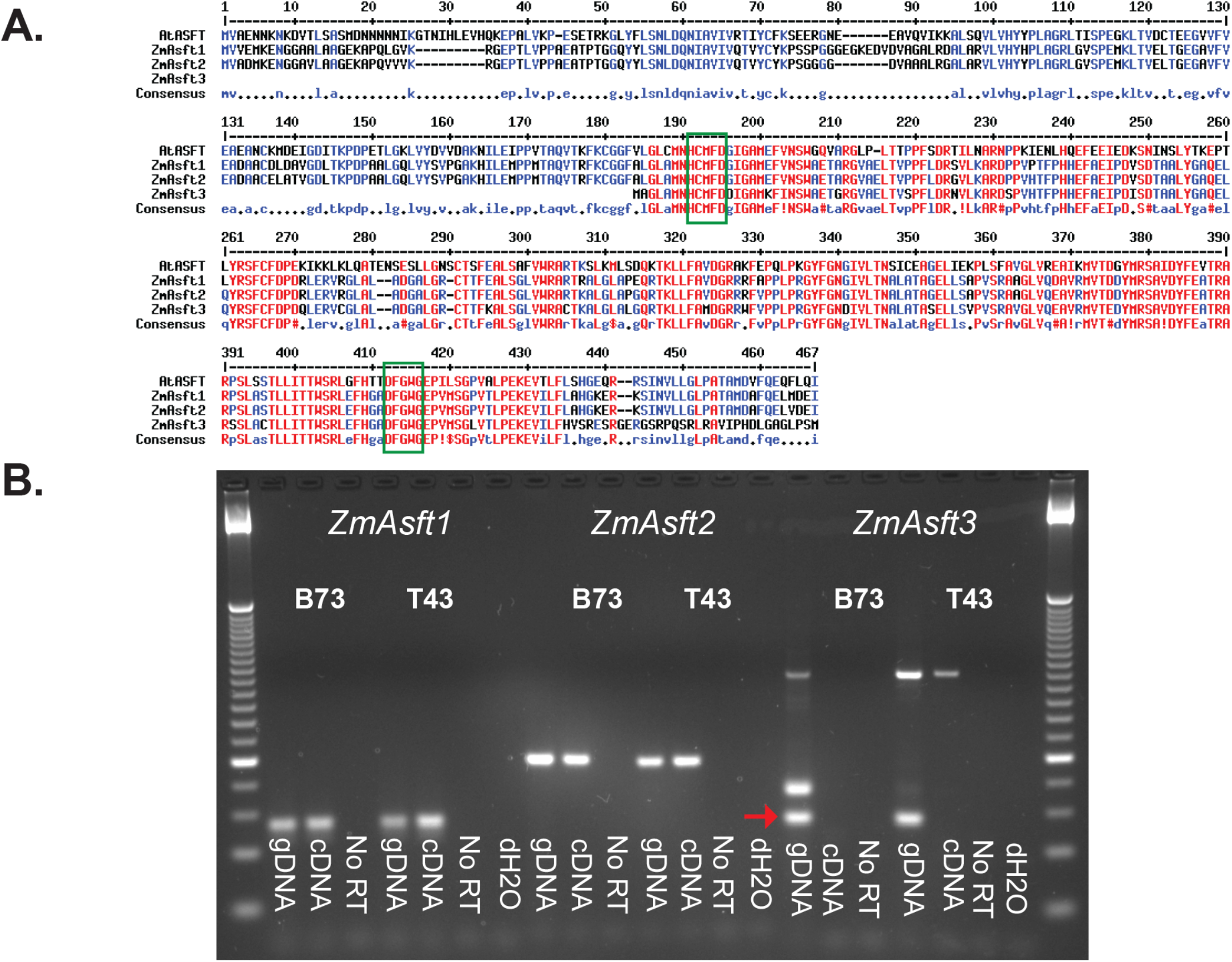
*ZmAsft3* is a putative pseudogene. **A.** Multiple sequence alignment of *ZmAsft* amino acid sequences with *AtASFT*. The canonical HXXXD and DFGWG motifs of BAHD acyltransferases are denoted by green boxes. **B.** Analysis of *ZmASFT* gene expression in developing leaves by RT-PCR. Gene-specific primers generate amplicons in both gDNA and cDNA samples, but not in cDNA samples prepared without reverse transcriptase (“No RT”) in both inbred backgrounds (B73 and T43 [W22]) for *ZmAsft1* and *ZmAsft2*. Conversely, the amplicon corresponding to the 3’UTR of *ZmAsft3* (red arrow) is absent from cDNA preparations.

**Supplemental Figure S4.**
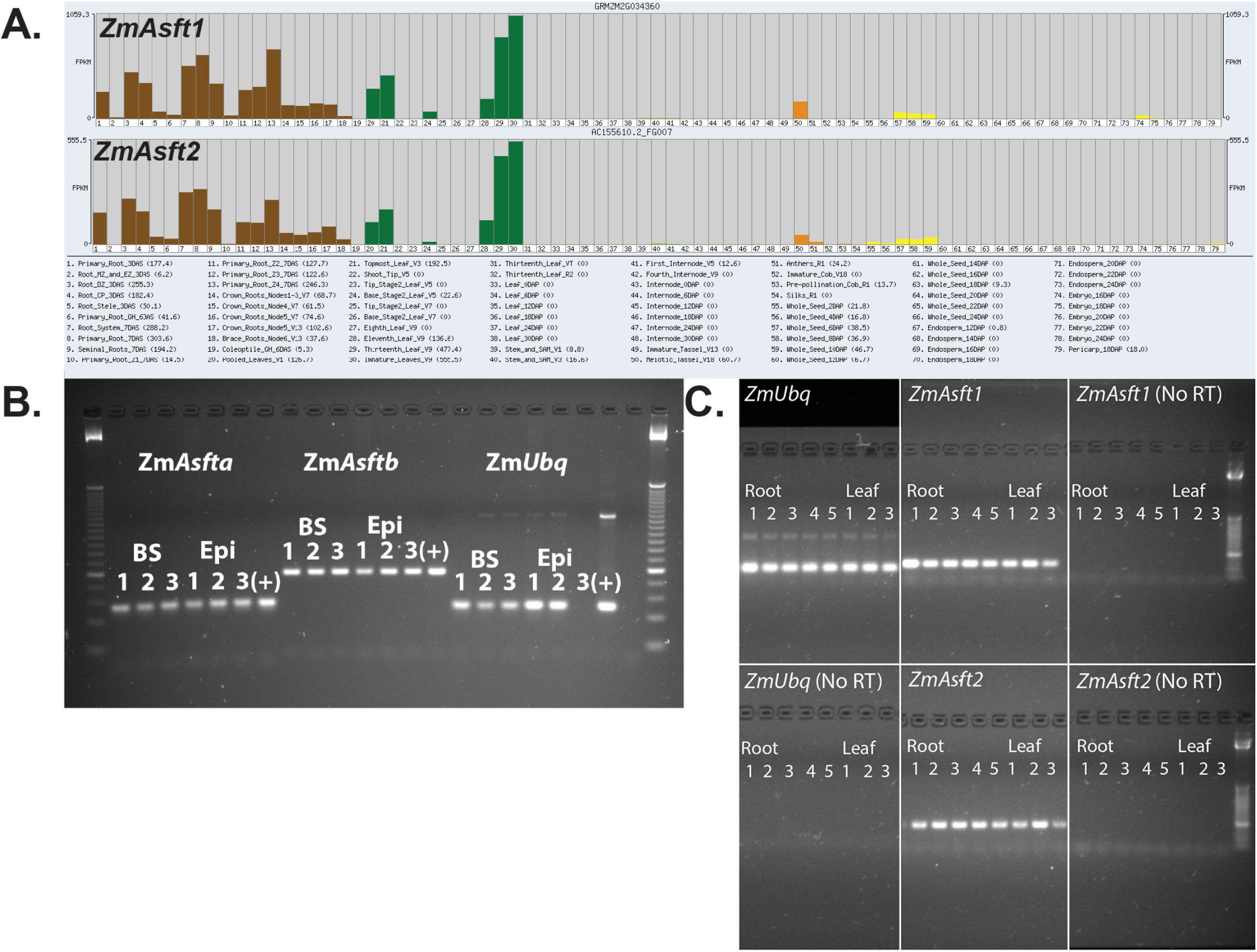
Organ and tissue-specific expression analysis of *ZmAsft* genes. **A.** RNA-Sequencing expression atlas of *ZmAsft1* and *ZmAsft2*. Gene records for *ZmAsft1* and *ZmAsft2* were accessed at Maize GDB (http://www.maizegdb.org), Both genes are most strongly expressed in developing primary roots (brown) and leaves (green). Expression is limited in tassels (orange) and developing seeds (yellow). **B.** RT-PCR of Laser-Capture Microdissected leaf tissues. *ZmAsft1, ZmAsft2*, and *ZmUbq1* expression was measured in triplicate samples of bundle sheath (BS) and epidermal cells (Epi), with pooled cDNA from undissected tissue serving as a control (+). The blank lane next to (+) contains a water blank. **C.** RT-PCR of developing seedling primary roots. Primary roots were divided into sections of equal length (Root 1-5; Section 1 is apical and Section 5 is basal). The basal 6 cm of developing leaves from the same plants were collected as a positive control (Leaf 1-3). *ZmAsft1, ZmAsft2*, and *ZmUbq* expression was evaluated as described in Supplemental Figure S3.

**Supplemental Figure S5.**
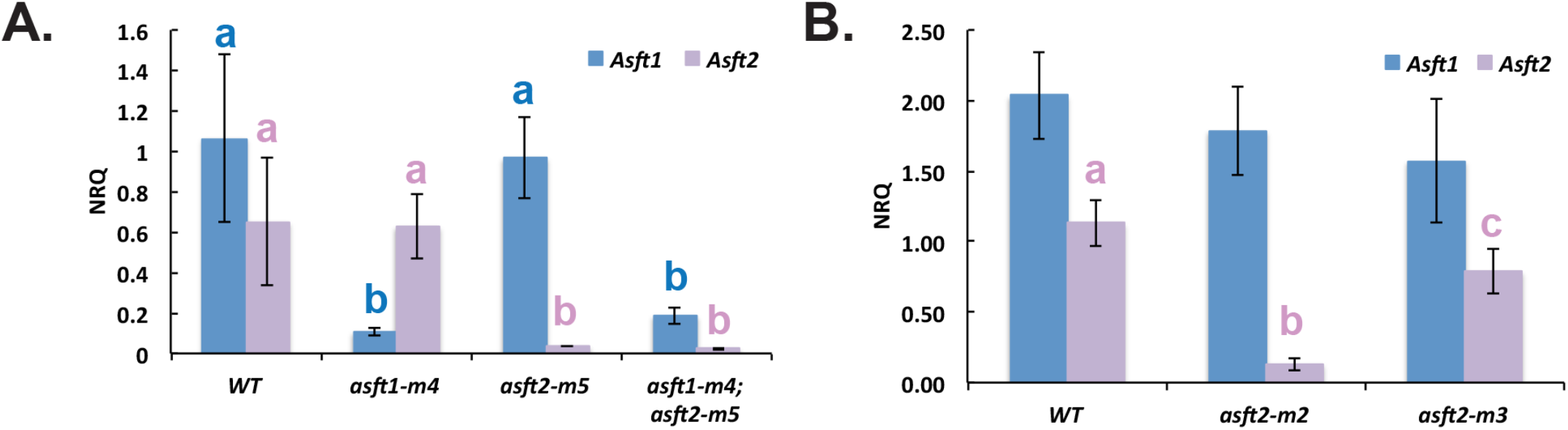
Expression analysis of additional single and double mutant alleles. **A.** qPCR analysis of *asft1-m4; asft2-m5* single and double mutants. **B.** qPCR analysis of *asft2-m2* and *asft2-m3* single mutants. Samples were analyzed as described in Figure 1C.

**Supplemental Figure S6.**
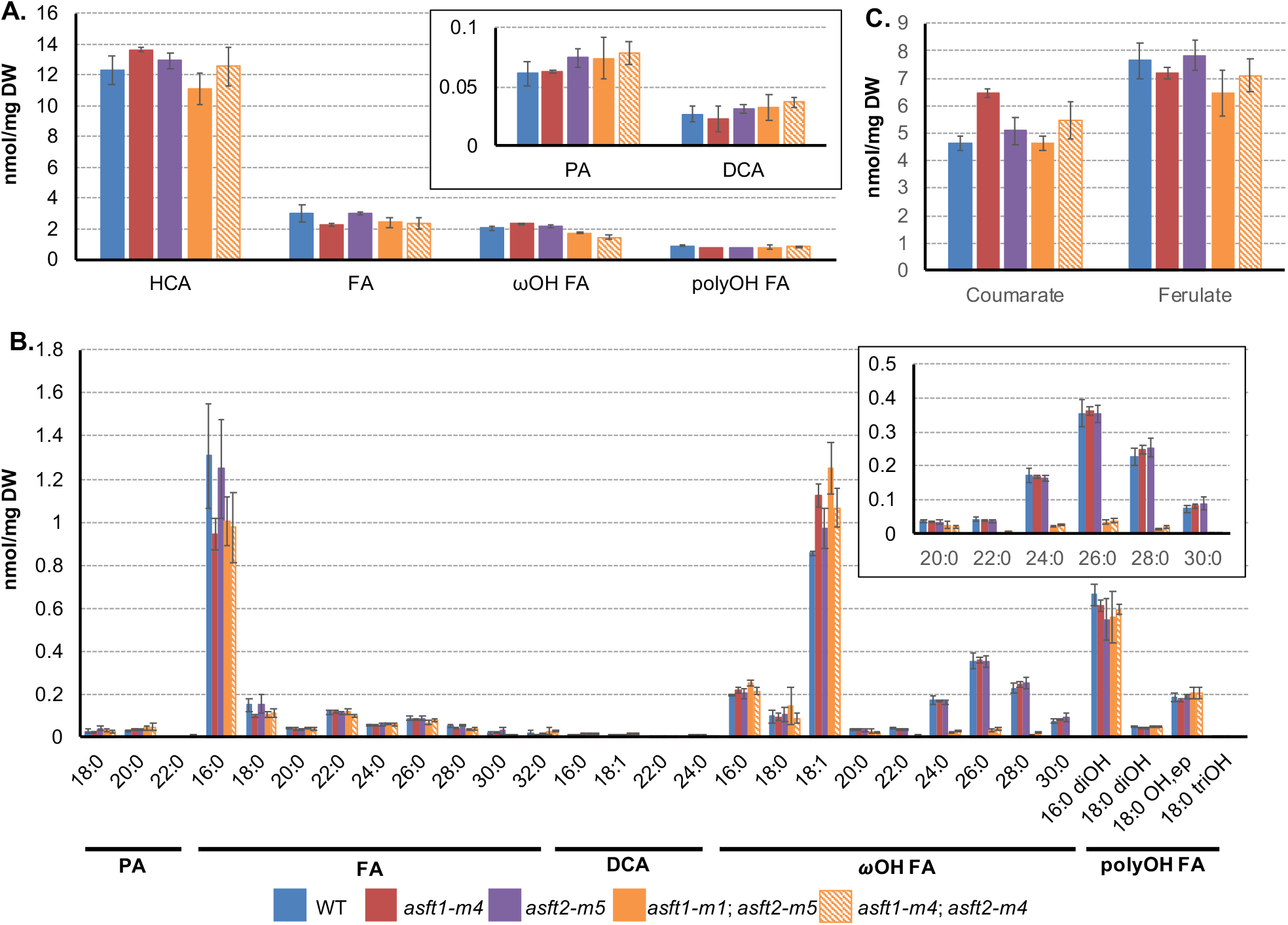
The aliphatic monomer defect is present in other double mutant combinations. **A.** Overview of the major monomer classes present in fully expanded third leaf laminae. Asterisks denote significant differences between genotypes determined by a one-way ANOVA (*,*p* < 0.05; *** p < 0.001) with a Tukey-Kramer post-hoc comparison (different letters denote significant differences at *p* < 0.05). Values are averages with standard deviations of three biological replicates. HCA, hydroxycinnamic acid; PA, primary alcohol; FA, fatty acid; DCA: α,ω-dicarboxylic acid; ωOH FA, ω-hydroxy fatty acid; polyOH FA, poly-hydroxy fatty acid. **B.** Aliphatic monomer content. Inset: Higher resolution image of C_20:0_-C_30:0_ ωOH FA. **C.** Hydroxycinnamic acid content.

**Supplemental Figure S7.**
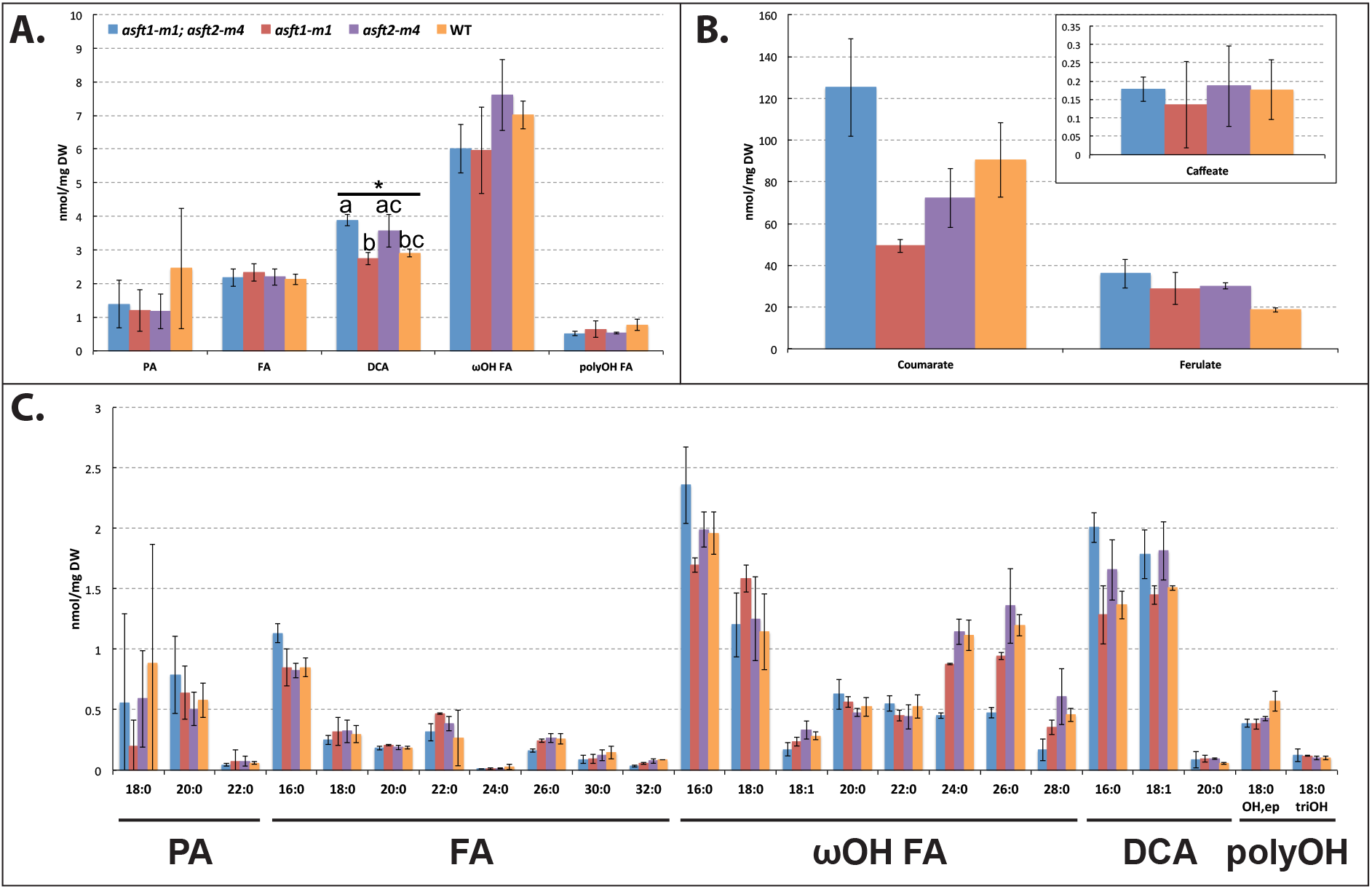
The root aliphatic monomer defect is specific to double mutants. **A.**Overview of the major monomer classes present in seedling primary roots. Values are averages with standard deviations of three biological replicates. HCA, hydroxycinnamic acid; PA, primary alcohol; FA, fatty acid; DCA: α,ω-dicarboxylic acids; ωOH FA, omega-hydroxy fatty acids; polyOH FA, poly-hydroxy fatty acids. B. Hydroxycinnamic acid content. **C.** Chain length distributions of aliphatic monomers. Horizontal axis labels denote chain lengths of individual monomers grouped by compound class (labels as described in **A.**).

**Supplemental Figure S8.**
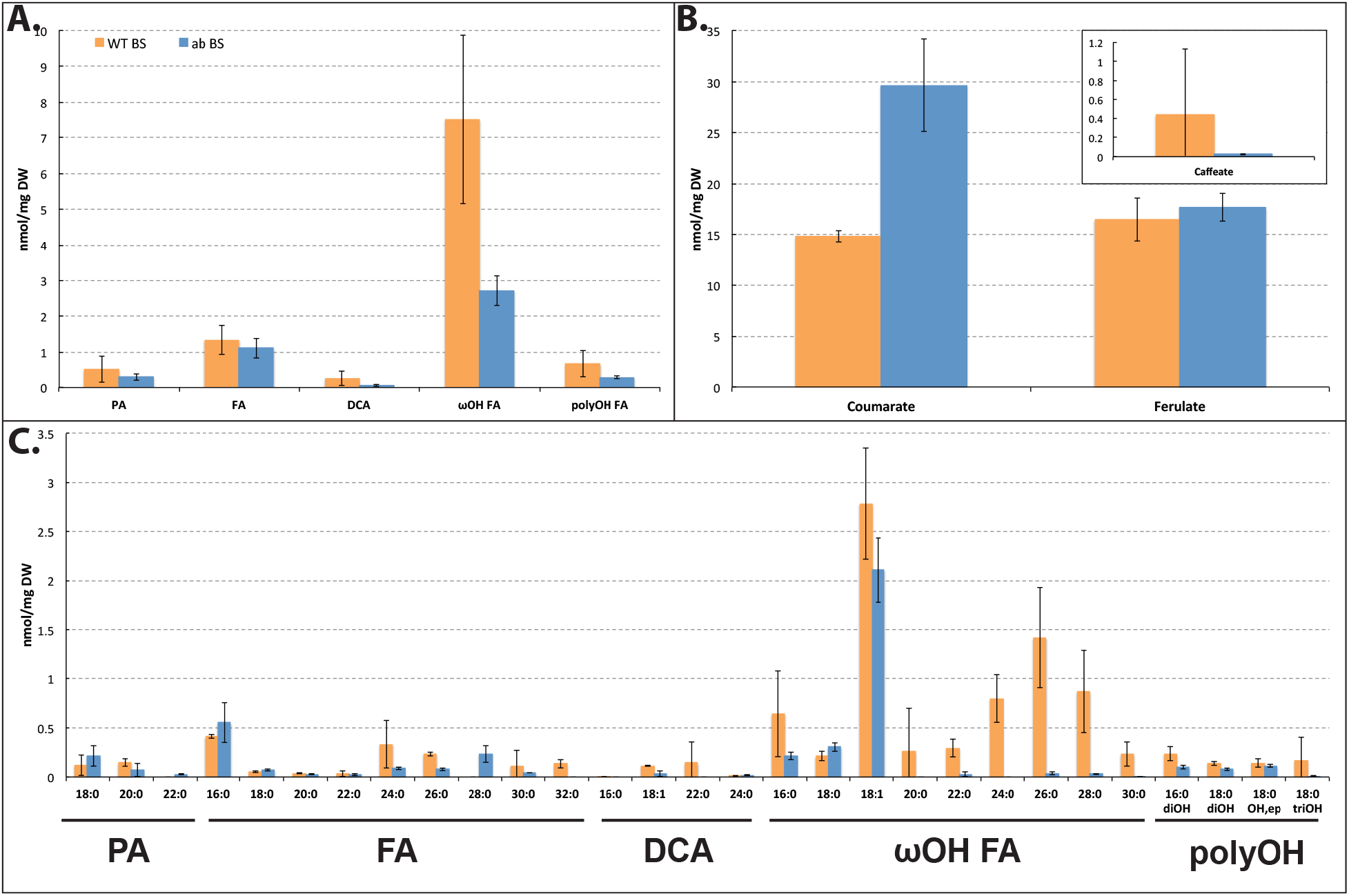
Polyester composition of purified bundle sheath strands. A. Overview of the major monomer classes present in WT *and asft1-m1; asft2-m4* third leaf laminae. Values are averages with standard deviations of three biological replicates. B. Hydroxycinnamic acid content of purified bundle sheath strands. C. Chain length distributions of aliphatic suberin monomers.

**Supplemental Figure S9.**
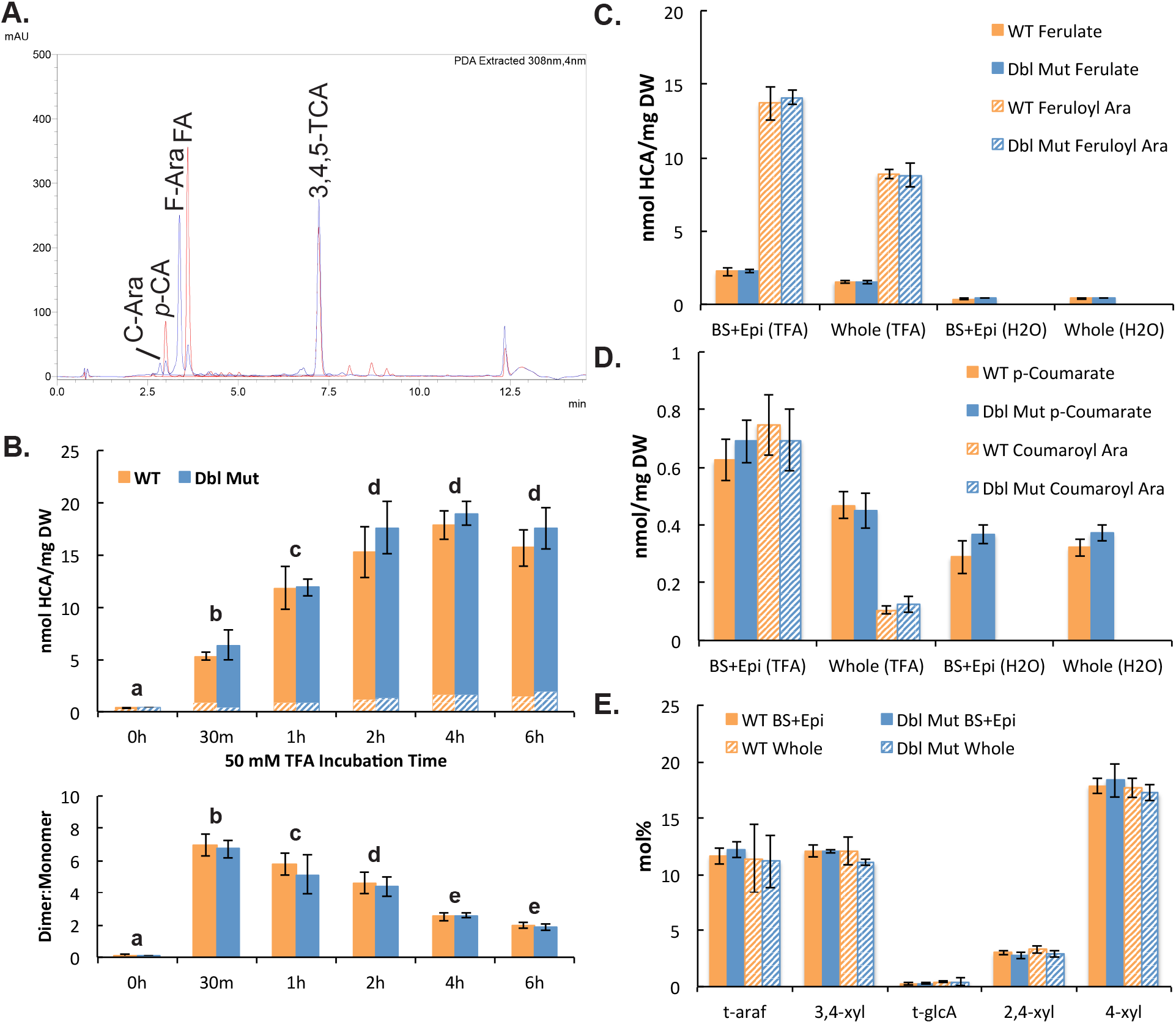
The majority of acid-labile ferulic acid in leaves is associated with glucuronoarabinoxylan. **A.** Representative UV-absorbance traces before and after saponification. UV absorbance (mAU) of TFA-soluble hydroxycinnamic acid (HCA) was analyzed by HPLC directly (blue trace) or after saponification to cleave alkali-labile ester linkages (red trace). Feruloyl arabinose (F-Ara) and *p*-coumaroyl arabinose (C-Ara) were converted to ferulic acid (FA) and p-coumaric acid (CA) by saponification. 100 μM 3,4,5-trimethoxy-*traws*-cinnamic acid (3,4,5-TCA) was introduced as an internal standard. **B.** Time course of HCA solubilization by TFA incubation. Lyophilized cell wall residues of wild type (WT, orange bars) and *asft1-m1*; *asft2-m4* double mutants (Dbl Mut, blue bars) were incubated in 50 mM TFA at 100°C for the indicated time (0h, 0.5h, 1h, 2h, 4h or 6h). Top panel: total ferulic acid (solid) and *p*-coumaric acid (hatched). Bottom panel: ratio of total dimers (feruloyl/coumaroyl arabinose) to monomers (ferulic/coumaric acid). Different letters indicate significant differences between treatments according to a Tukey-Kramer post-hoc test (*p* < 0.05) following a significant two-way ANOVA (treatment x genotype, treatment effect *p* < 0.001). There was no genotype effect. Values are averages plus standard deviations of three biological replicates per treatment. **C.** Acid-labile ferulic acid derivatives from fractionated (BS+Epi) and whole leaves incubated at 100°C for 2h in 50 mM TFA or water. There was no significant difference in feruloyl arabinose dimers or ferulic acid monomers between genotypes for either cell wall preparation (one-way ANOVA, *p* > 0.05). Values are averages plus standard deviations of six biological replicates. **D.** Acid-labile *p*-coumaric acid derivatives from fractionated (BS+Epi) and whole leaves. Sample preparation and statistical analysis are described in Figure 4C. **E.** Linkage-methylation analysis of monosaccharide linkage groups from glucuronoarabinoxylan. The polymer consists of a linear β-(1à4)-D-xylose backbone (4-xyl) decorated with terminal (5-O-feruloyl/coumaroyl) α-L-arabinofuranose-(1 à3)-β-D-xylose branches (*t*-araf, 3,4-xyl). The polymer also contains a small number of terminal α-D-glucuronic acid-(1 à2)-β-D-xylose branches (*t*-glcA, 2,4-xyl). None of the linkages are significantly different between genotypes for either cell wall preparation (one-way ANOVA, *p* > 0.05). Values are averages with standard deviations for two biological replicates with two technical replicates each.

**Supplemental Figure S10.**
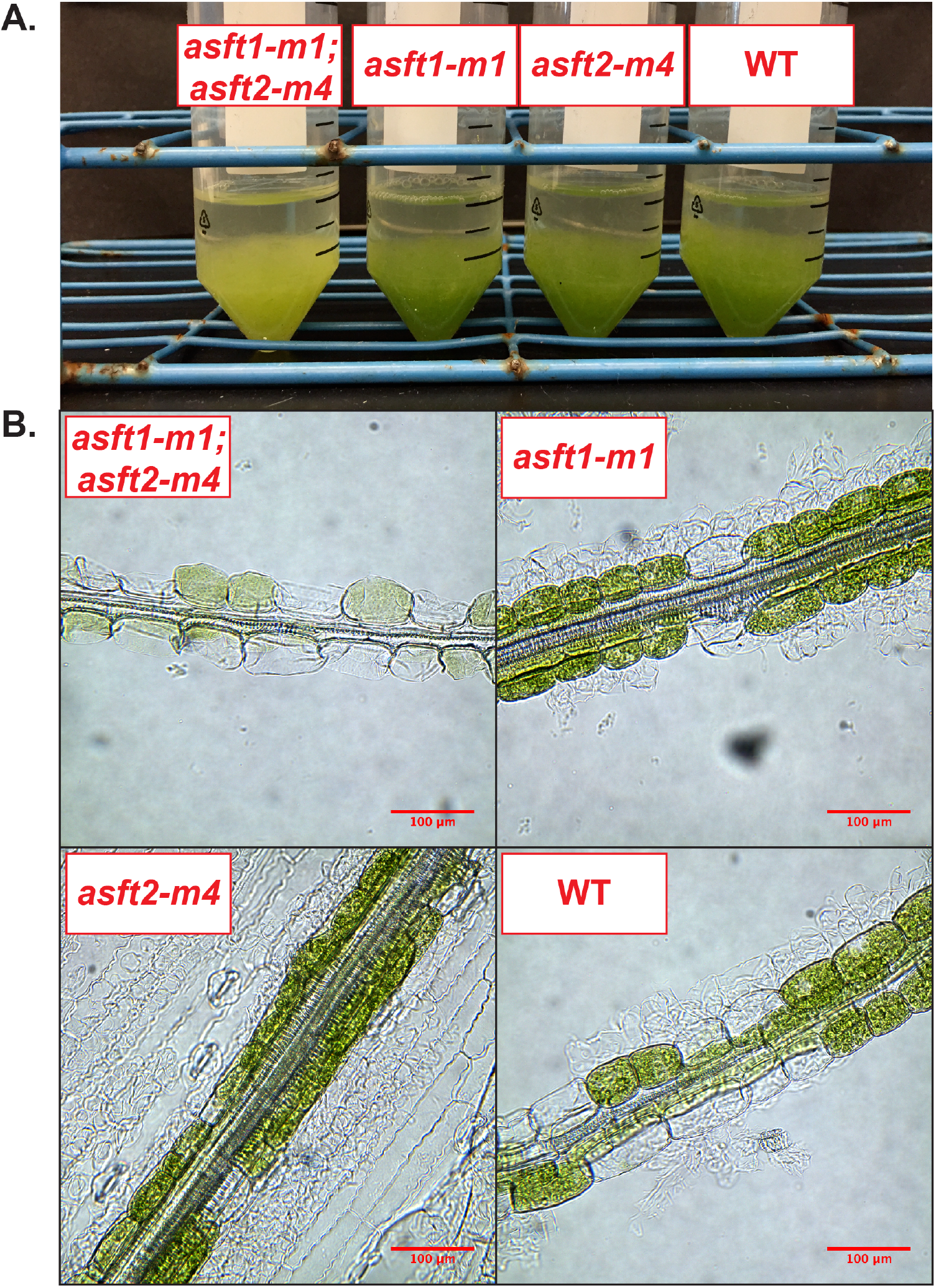
Double mutant bundle sheath cell walls are fragile relative to wild type. **A.** Homogenate of bundle sheath strands and epidermal cells from homogenized third leaf laminae of wild type (WT) and *asft* mutants. **B.** Bright field microscope images of sheared bundle sheath strands. Nearly all of the bundle sheath cells of the double mutant are broken, and the chlorophyll has leaked out during sample preparation. Single mutant and WT bundle sheath strands retain many intact cells with abundant chlorophyll. A segment of intact epidermis with pavement cells and stomata is visible in the *asft2-m4* image. 200x magnification. Scale bars denote 100 μm.

**Supplemental Figure S11.**
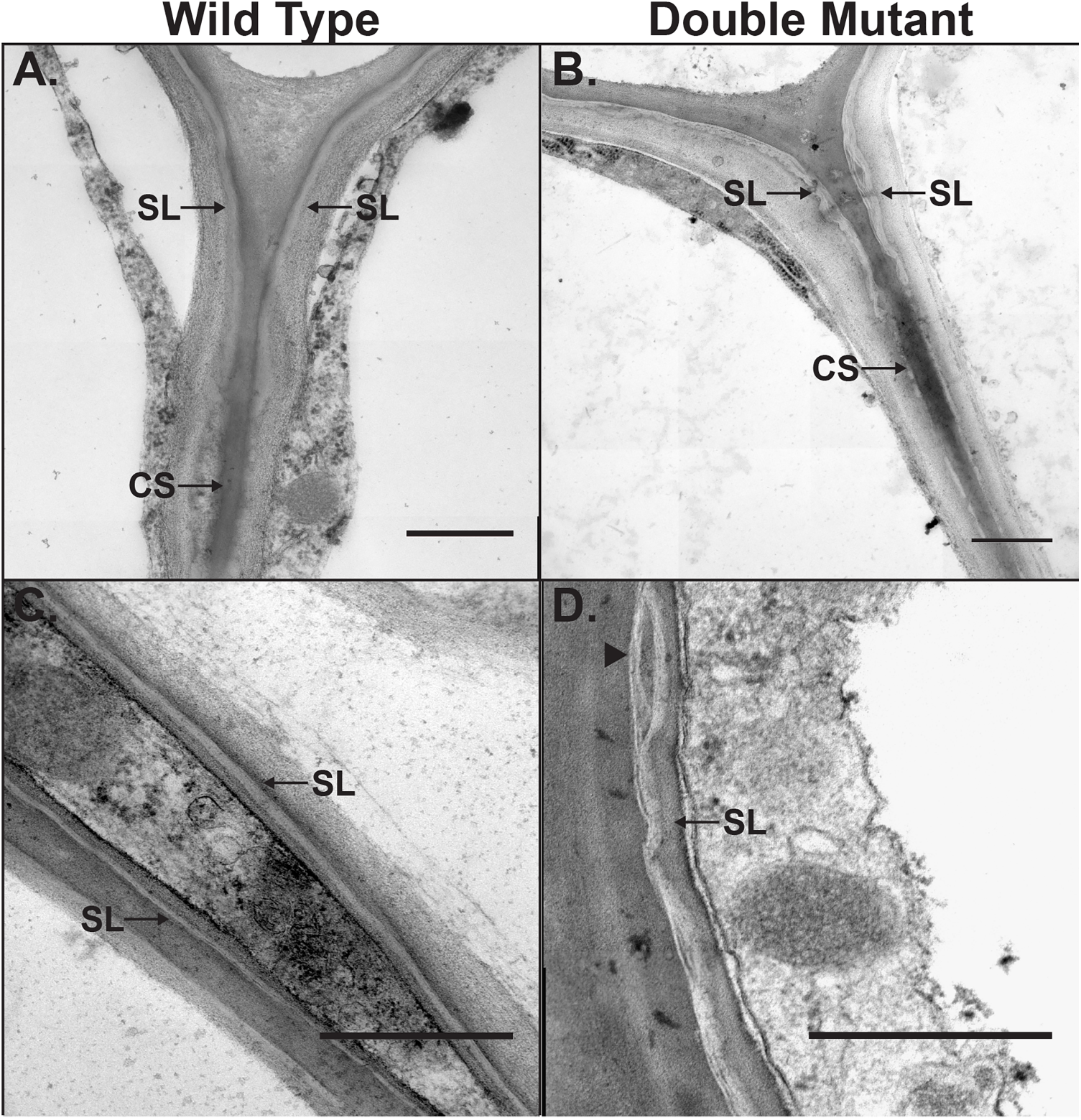
The *ZmAsft* genes are required for normal suberin lamella ultrastructure in the hypodermis. **A.** Transmission electron micrograph of inner tangential and radial cell walls of the wild type endodermis near a cell corner. Suberin lamellae (SL) and Casparian Strips (CS) are denoted by arrows. 25,000x magnification. Scale bar denotes 200 nm. **B.** Inner tangential and radial cell walls of the double mutant endodermis near a cell corner. 25,000x magnification. Scale bar denotes 200 nm. **C.** Suberized radial cell walls of two adjacent hypodermal cells from wild type. The SL exhibit a uniform “tramline” architecture. 16,000x magnification. Scale bar denotes 200 nm. **D.** Suberized outer tangential cell wall of the double mutant hypodermis adjacent to an epidermal cell. The SL are electron lucent and irregular in thickness compared to WT. Occasional inclusions of electron material of comparable texture to the adjacent polysaccharide cell wall are apparent (arrowhead). 20,000x magnification. Scale bar denotes 200 nm.

**Supplemental Table S1.**
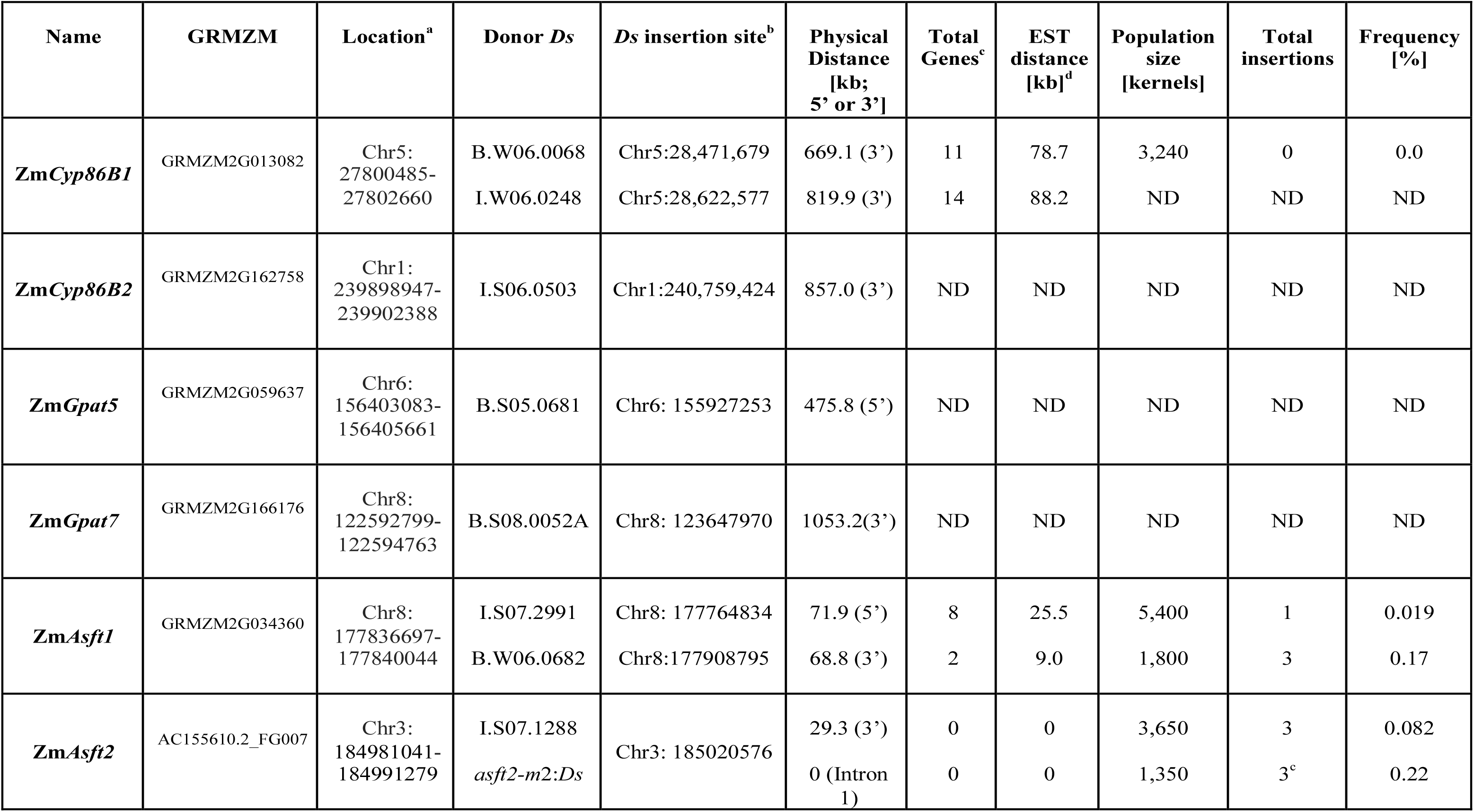
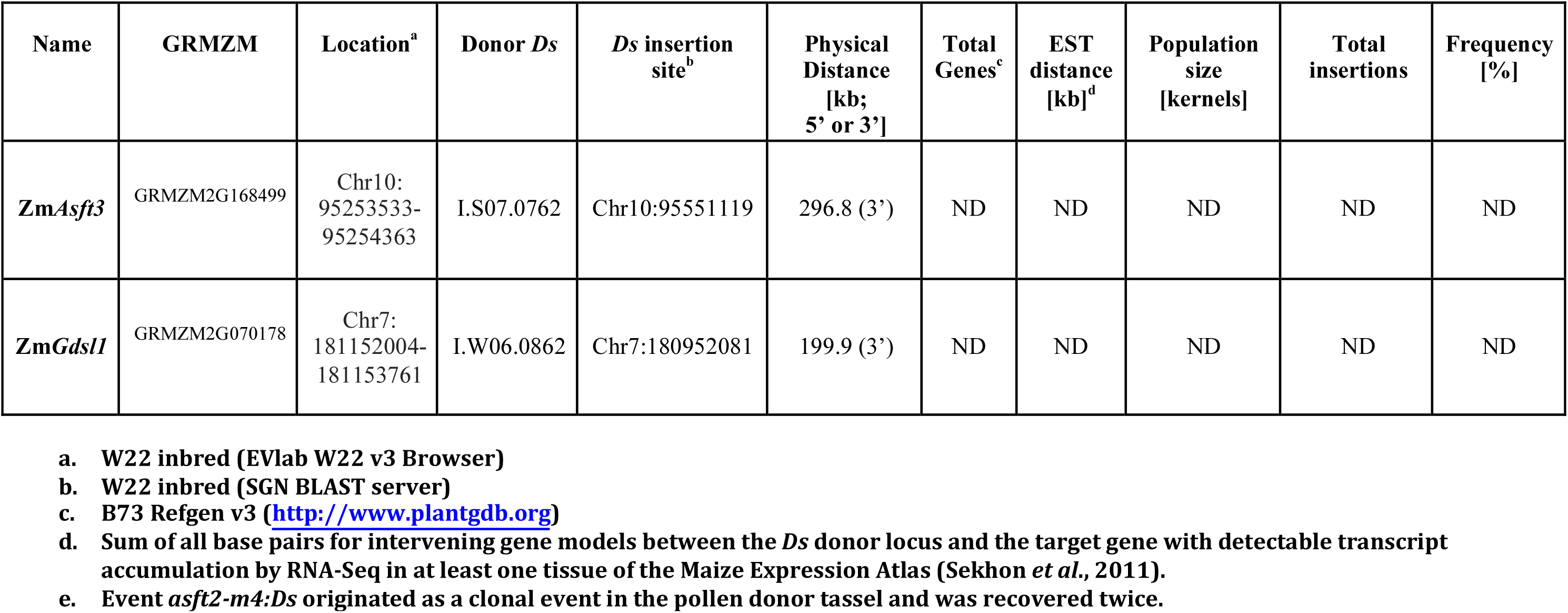
Targeted mutagenesis of suberin biosynthesis candidates using *Dissociator (Ds)* transposons.

**Supplemental Table S2.**
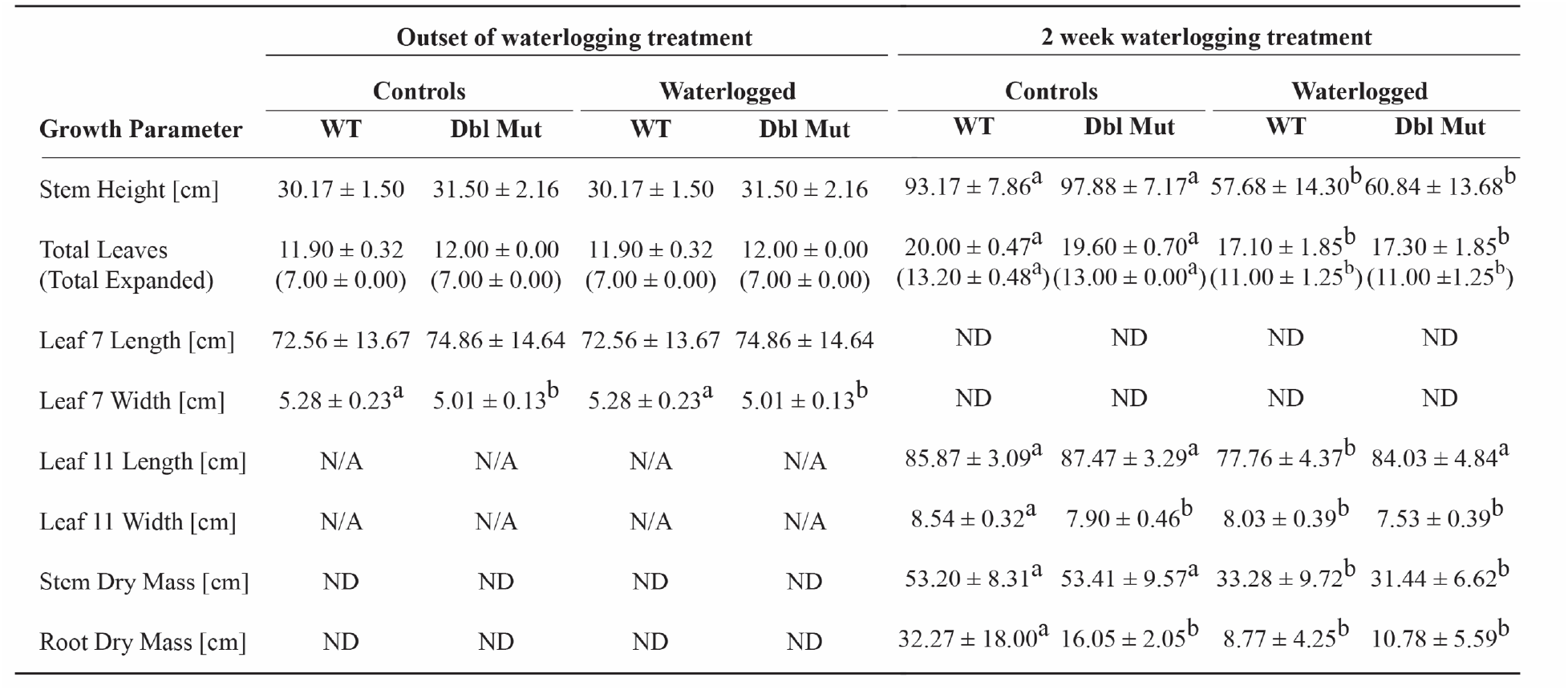
Table 1. Growth of wild type and double mutant plants during waterlogging stress.

**Supplemental Table S3.**
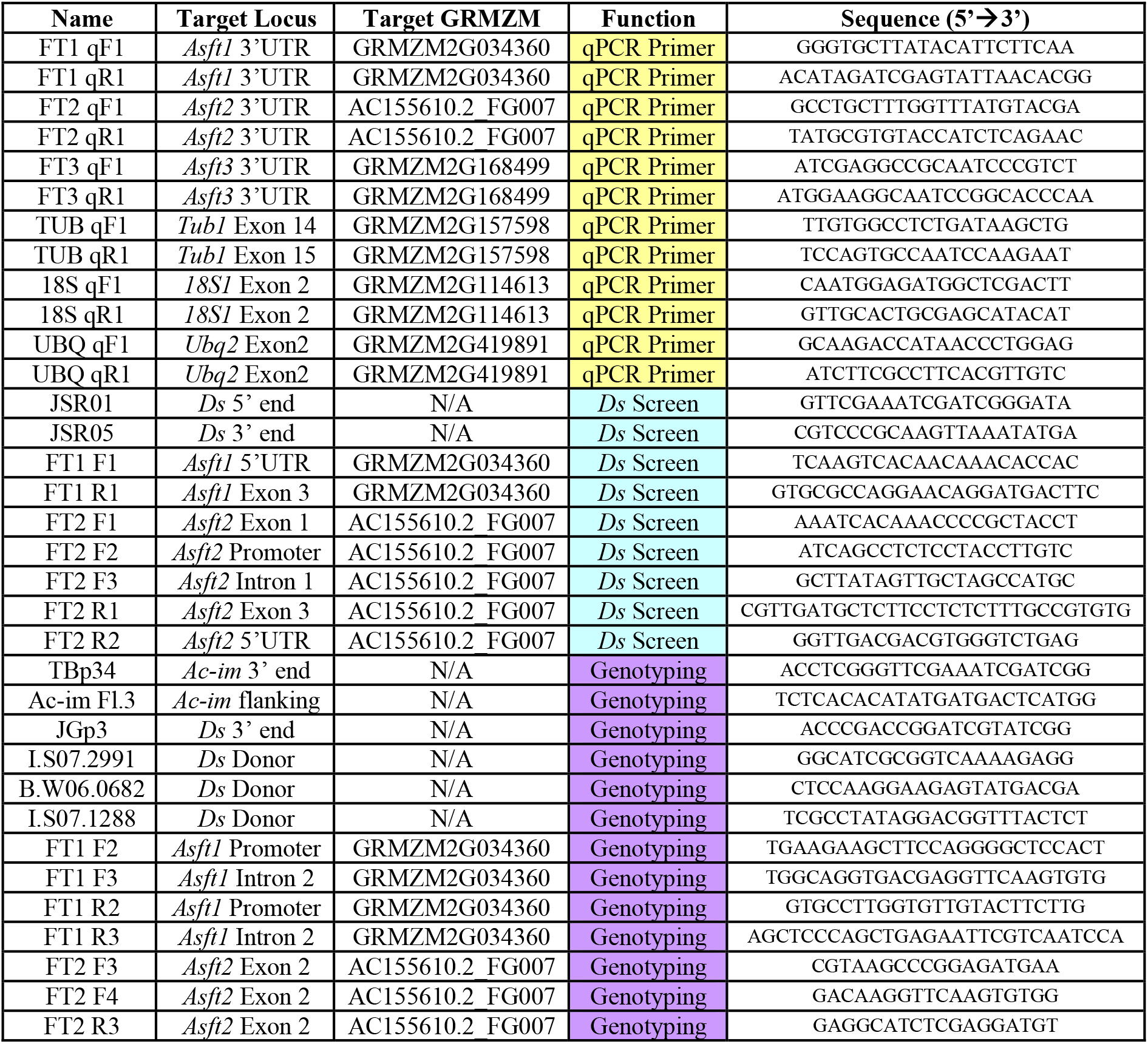
Supplemental Table S2 Primers used in this study.

**Supplemental Table S4.**
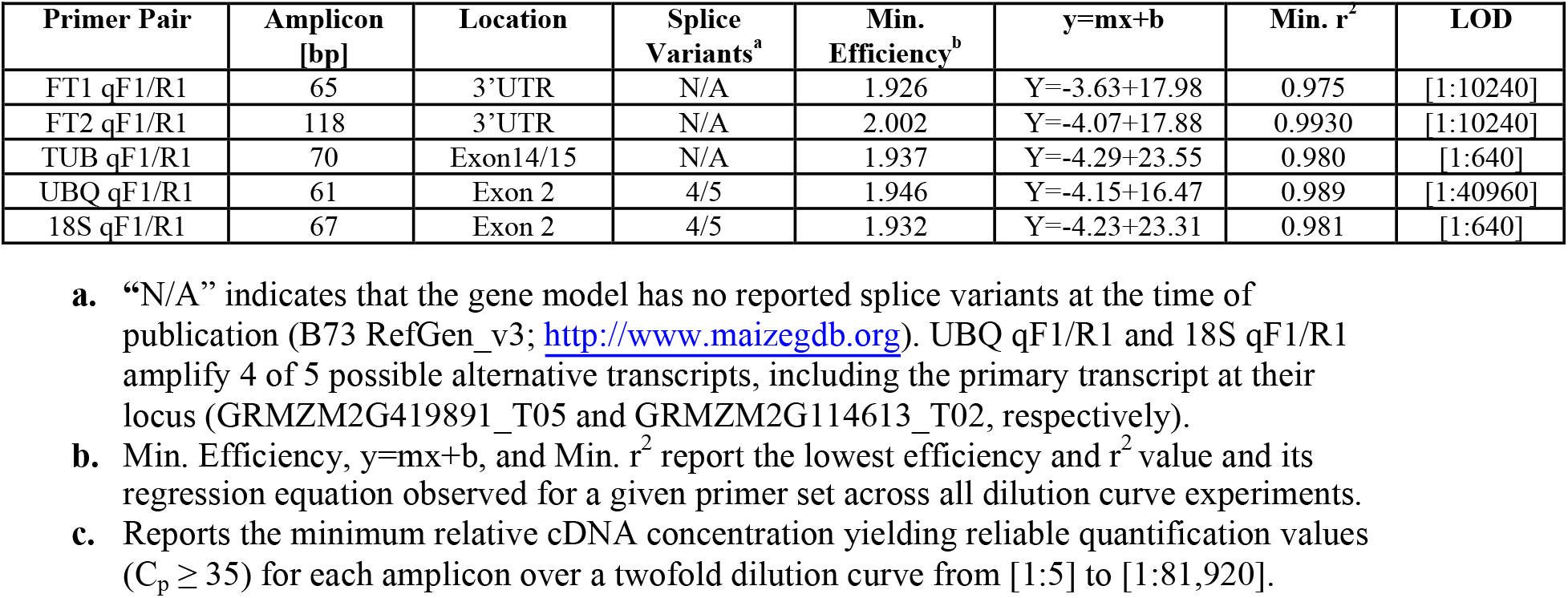
Supplemental Table S3. Quality control metrics for qPCR primers.

