## Supplemental Methods for "The maize Aliphatic Suberin Feruloyl Transferase genes affect leaf water movement but are dispensable for bundle sheath CO_2_ concentration"

**Phylogenetic analysis of candidate *Asft* genes**

The *Arabidopsis thaliana* ALIPHATIC SUBERIN FERULOYL TRANSFERASE amino acid sequence was queried against the maize (*Zea mays*; B73 RefGen_v3), sorghum (*Sorghum bicolor* v3.1), switchgrass (*Panicum virgatum* AP13 v1.0), foxtail millet (*Setaria italica* v2.2), green millet (*Setaria viridis* A10.1 v1.1), rice (*Oryza sativa* Nipponbare/*japonica* v7.0), and Brachypodium (*Brachypodium distachyon* Bd21 v3.1) reference proteomes using BLAST-P searches under the default parameters at Phytozome 11 (<http://www.phytozome.net>; Goodstein *et al*., 2012). Previously characterized amino acid sequences from model dicots (AtASFT, AtDCF, AtFACT, PtHHT, and StFHT) were retrieved from Phytozome 11, and all amino acid sequences were aligned within the MEGA environment using MUSCLE under the default parameters. A Neighbor-Joining tree was generated from the alignments using a Jones-Taylor-Thornton model assuming uniform substitution rates, and pairwise deletion of gaps. Branch support was estimated with 1000 bootstrap replications. The ASFT sub-clade formed a monophyletic group within Clade V of the BAHD acyltransferases (Tuominen *et al*., 2011). AtHCT, a BAHD acyltransferase involved in lignin biosynthesis (Hoffmann *et al*., 2004), served as the outgroup.

**Cell wall HCA fractionation**

Delipidated cell wall residues were incubated overnight in 90% DMSO on a rocking agitator (Carpita, 1983), pelleted, and washed twice with MilliQ-grade water. Samples were flash-frozen and lyophilized.

For HCA fractionation, the method of Saulnier *et al*. (1995) was utilized. 20 mg lyophilized material plus 100μM 3,4,5- trimethoxy-*trans*-cinnamic acid as an internal standard was incubated at 100°C in 3 mL of 50 mM TFA or water (negative control). After the designated incubation period (0.5, 1, 2, 4, or 6 hours), residual solids were pelleted and the supernatant was divided equally. Both aliquots were mixed with 0.5 mL *tert*-butanol and air-dried overnight. The first aliquot was solubilized with 50% (v/v) methanol and analyzed immediately by reverse-phase HPLC as described below. The second aliquot was saponified with 1 mL of 1M NaOH for 30 minutes in a 42 °C water bath, acidified with 1 mL of 3M HCl, and extracted three times with ethyl acetate. The combined ethyl acetate extracts were dried under nitrogen and solubilized in 50% methanol for HPLC.

HPLC was carried out on a Dionex Ultimate 3000 HPLC system (Thermo Scientific-Dionex) with UV detection (320 nm) using an SPD-M20A photodiode array detector (Shimadzu). Samples were partitioned on a reverse-phase C18 (Shimadzu Shim-pack XR-ODS, 3 mm i.d. x 0.75 mm length 2.2 μm bead diameter) column maintained at 40**°**C. Solvents A (0.1% v/v formic acid) and B (100% acetonitrile) were supplied at a flow rate of 0.7 mL/min according to the following program: 0 to 0:30, 5% B isocratic; 0:30 to 0:40 5% to 10% B linear; 0:40 to 11:00, 10% to 25% B linear; 11:00 to 11:20, 25% B to 95% B linear; 11:20 to 12:20, 95% B isocratic; 12:20 to 13:10, 95% to 5% B linear; 13:10 to 14:00, 5% B isocratic.

**Linkage-methylation analysis**

Linkage-methylation analyses were carried out as described in Mertz *et al*. (2012). Duplicate samples from two biological replicates were prepared for mechanically fractionated and whole leaf samples of WT and double mutants. 10 mg of lyophilized residue per sample was carboxyl-reduced with NaBD_4_ in the presence of a water-soluble diimide as described in Kim and Carpita (1992) and Carpita and McCann (1996). Samples were dialyzed against running water for 48 hours, frozen, and lyophilized. Pellets were partitioned into three 1-2 mg samples; the first sample was prepared for monosaccharide compositional analysis using the alditol acetates protocol as described in Gibeaut and Carpita (1991). A linkage-methylation analysis was conducted from the second and third sample as technical duplicates according to Gibeaut and Carpita (1991).

**Waterlogging Stress Experiment**

The waterlogging stress experiment was modeled after Abiko *et al*. (2012). At 28d after sowing under ambient greenhouse conditions, double mutant and WT individuals were divided into control and treatment groups (n = 10 plants per genotype per treatment). Plants assigned to the waterlogging treatment group were submerged in a hydroponic box (8’ L x 4’ W x 1.5’ H) containing stagnant water maintained at a depth of 2 cm above the soil line. WT and double mutant plants were arranged in an alternating pattern to minimize shading and positional effects. Control plants were arranged in randomized blocks adjacent to the submergence box. After two weeks, plant height, total leaves, dimensions of the youngest fully expanded leaf common to both treatment groups, and shoot and root dry weight were recorded. Genotype and treatment effects were evaluated by two-way ANOVA [treatment x genotype] with a Bonferroni-Holm multiple testing correction. A Tukey’s HSD post-hoc test was conducted at a 95% significance level following a significant ANOVA result.
